# Type-I interferon priming signal balances anti-bacterial and anti-tumor trained immunity in alveolar macrophages

**DOI:** 10.1101/2025.06.14.659659

**Authors:** Tao Wang, Jinjing Zhang, Yuxuan Miao, Yanling Wang, Ying Li, Lu Wang, Yuanyuan Liu, Chia-Wei Chang, Enguo Chen, Yushi Yao

## Abstract

Tissue-resident macrophages may be trained to confer enhanced response to heterologous re-stimulation, and thus foster versatile trained immunity (TI) against both infections and tumors. However, key priming signals that contribute to such functional versatility in trained macrophages remain not fully understood. Here we show that influenza A virus (IAV) infection in mice induces lasting transcriptional imprints of acute type-I interferon (IFN-I) signaling in lung-resident alveolar macrophages (AMs) that confer balanced anti-bacterial and anti-tumor TI responses. In both IAV-infected mice and an *ex vivo* cytokine-mediated training system, IFN-I signaling is found to be critical to developing anti-tumor TI functions, although it limits anti-bacterial TI functions in AMs. Moreover, human AMs carrying transcriptional imprints of IFN-I signaling are associated with anti-tumor immunity and immune activation in lung cancer tissues. Our findings highlight IFN-I as a key priming signal required to developing versatile TI functions in AMs with balanced anti-bacterial and anti-tumor potential.

## Introduction

Trained immunity (TI) represents *de facto* innate immune memory induced by a prior stimulation in innate leukocytes to foster their potential of enhanced responsiveness to a broad range of heterologous secondary stimulation^1–3^. Development of TI/innate immune memory was shown to be primed by microbial components and/or proinflammatory cytokines that induce prolonged epigenetic and metabolic reprogramming, resulting in long-lasting transcriptional, phenotypic and functional changes in trained/memory innate leukocytes^1,3–5^. A key difference between TI and adaptive immune memory is that the former is not strictly antigen-specific due to the lacking of antigen-specific receptors in innate leukocytes^3^. Therefore, trained innate leukocytes can respond to a broad spectrum of secondary stimulants, and TI has been associated with versatile host defense functions including enhanced anti-bacterial and anti-tumor innate immunity in both experimental and clinical scenarios^6–14^. In this regard, microbes and microbial components including but not limited to BCG, β-glucan, and respiratory viruses were shown to induce both anti-bacterial and anti-tumor TI in innate myeloid cells^8,9,11,12,14–16^. Although a few key molecular pathways, such as IFN-γ, IL-1β, or dectin-1, were respectively identified as a single key signal required to the priming of TI in various infections^12,17,18^, the mechanisms through which such versatile (i.e., anti-bacterial and anti-tumor) TI functions are induced have remained not fully understood.

TI-priming stimulation (e.g., microbial infection) typically activates multiple innate and adaptive immune pathways and therefore induce a milieu of proinflammatory cytokines. It remains unclear whether 1) the versatile (i.e., anti-bacterial and anti-tumor) TI functions are acquired by innate leukocytes as an integrated and inseparable package induced by one single priming signal (e.g., IFN-γ), or 2) various priming “ingredients” (e.g., multiple proinflammatory cytokines and/or ligands of pattern recognition receptors) respectively shape different functional aspects of TI and therefore jointly induce versatile TI functions. These unanswered key questions are highly relevant to both understanding the mechanisms of TI development and the potential clinical application of TI which may require disease-specific tailoring of TI potentially via introducing a formular of priming stimulants.

Alveolar macrophages (AMs) represent a lung tissue-resident macrophage subset that play critical roles in host defense against pulmonary diseases including bacterial pneumonia and lung cancer^19–21^. Derived originally from embryonic hematopoiesis, AMs are self-sustaining in steady state and even after respiratory inflammation^11,12,19,22^. Therefore, AMs can carry long-lasting and tissue-specific innate immune memory independently of “systemic/central TI” in circulating monocytes or bone marrow myeloid progenitors^11,12,23^. Current experimental evidence suggests that respiratory infection-induced TI in AMs is a key factor that shapes their long-term host defense functions against respiratory infections and lung cancer^11,12,16^. Specifically, we have shown in previous studies that respiratory viral infection-triggered IFN-γ priming signal is required to the development of both anti-bacterial and anti-tumor TI functions in AMs^11,12^. IFN-γ was also shown to be required to *Cryptococcus neoformans*-induced TI in lung macrophages with enhanced defense functions against homologous re-infection^24^. Beyond the critical roles of IFN-γ, it remains largely unclear whether and how other pro-inflammatory cytokines contribute to the development and in particular the functional versatility of TI in AMs.

Type I interferon (IFN-I) represents a characteristic proinflammatory cytokine abundantly produced early after respiratory viral infections such as influenza A virus (IAV) infection. IFN-I is produced by a variety of leukocytes and non-hematopoietic cells, and receptor of IFN-I is ubiquitously expressed^25^. Not unexpectedly, IFN-I was shown to modulate the functions of a wide variety of cell types including AMs^23,26^. Specifically, IAV-induced IFN-I was shown to acutely suppress the anti-bacterial functions of AMs and therefore contributes to secondary bacterial super-infection^26^. Moreover, IFN-I was shown to induce long-lasting epigenetic changes in multiple cell types including macrophages, suggesting its potential in shaping long-term immune functions of macrophages^27^. Indeed, a more recent study showed that IFN-I was required to LPS-induced TI development in AMs^23^. Despite these intriguing and important findings, the potential roles of IFN-I in regulating the anti-bacterial versus anti-tumor TI functions in AMs remain largely unappreciated.

Here, we show in a mouse lung metastatic tumor model introduced long after acute respiratory IAV infection that IAV-induced acute IFN-I signaling is key to developing anti-tumor TI functions in AMs. Such a critical role of IFN-I in priming anti-tumor TI functions in AMs was also evidenced in an *ex vivo* cytokine-mediated AM training system with IFN-I and/or IFN-γ. Intriguingly, we show a contrasting role of IFN-I on the development of anti-bacterial TI functions in both mouse *in vivo* experiments and *ex vivo* AM training system, where IFN-I limits anti-bacterial functions in trained AMs. These data collectively suggest that, unlike IFN-γ that is required to inducing both anti-bacterial and anti-tumor TI functions in AMs, IFN-I signaling works in synergy with IFN-γ and preferentially induces anti-tumor TI functions while IFN-I limits IFN-γ-induced anti-bacterial TI functions in AMs. Therefore, IFN-I priming signal is key to the functional versatility of TI via balancing anti-bacterial and anti-tumor functions in trained AMs.

## Results

### IAV-trained AMs confer enhanced response to both bacteria and tumor cells

To determine the responsiveness of trained AMs to secondary bacterial and tumoral stimulations, wild type (WT) C57BL/6 mice were infected intranasally (i.n.) with a sublethal dose of IAV (A/PR8/34 strain) to induce TI in AMs^12^. Purified AMs from uninfected or IAV-infected mice at 30 days post infection were stimulated *ex vivo* with lipopolysaccharide (LPS), live *Streptococcus pneumoniae* (S.P.), or B16 melanoma cells, followed by quantification of selected pro-inflammatory cyto-/chemokines in cell culture supernatants and analysis on transcriptional changes in AMs (Figure 1A). IAV-trained AMs, in response to live S.P. or LPS, secreted higher amount of proinflammatory cyto/chemokines including TNF, IL-6, and macrophage inflammatory protein 2 (MIP-2) but not IFN-γ than AMs from uninfected (PBS) mice (Figure 1B). RNA sequencing (RNA-seq) analysis in purified AMs showed that, after S.P. stimulation, there were drastic transcriptional changes in both PBS-AMs (transcription of 12820 genes upregulated and 5067 downregulated) and IAV-AMs (8022 genes upregulated and 4450 downregulated) (Figures 1C, S1A, and S1B). Further analysis on differentially expressed genes showed that S.P. induced more immune function-related gene transcripts in IAV- than PBS-AMs, including genes related to TNF and IL-6 production, pattern recognition receptor signaling pathway, and response to molecule of bacterial origin (Figures 1D, 1E, and S1C). These data suggest that IAV-trained AMs are hyperresponsive to bacteria both functionally and transcriptionally.

**Figure 1.**
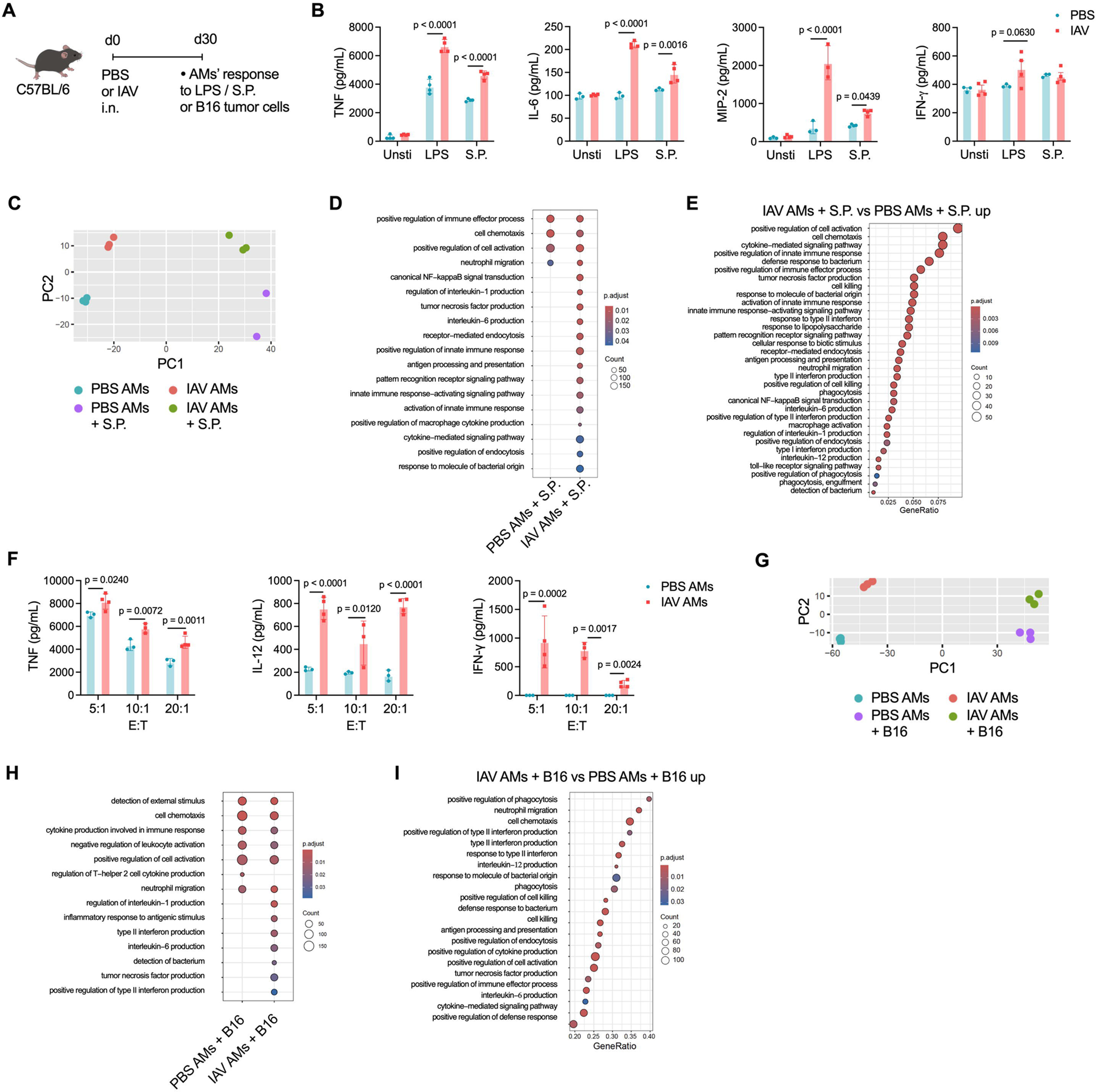
IAV-trained AMs confer enhanced response to both bacteria and tumor cells. (A) Schema of intranasal (i.n.) influenza A virus (IAV) infection in mice and *ex vivo* re-stimulation of purified AMs on day 30 after IAV infection. (B) Concentrations of representative proinflammatory cyto-/chemokines, including TNF, IL-6, MIP-2 and IFN-γ, in supernatants of cultured AMs isolated from IAV-infected (IAV AMs) or uninfected (PBS AMs) mice. AMs were either unstimulated (Unsti) or stimulated *ex vivo* with LPS or live *Streptococcal pneumoniae* (S.P.). (C) PCA plot of gene expression in S.P.-stimulated or unstimulated AMs. (D) GO enrichment of gene transcripts upregulated in PBS or IAV AMs upon S.P. stimulation as compared to unstimulated cells. (E) Go enrichment of gene transcripts upregulated in S.P.-stimulated IAV versus PBS AMs. (F) Concentrations of representative proinflammatory cytokines, including TNF, IL-12 and IFN-γ, in supernatants of PBS or IAV AMs cocultured for 72 hours with B16 melanoma cells at different E:T (AMs versus tumor cells) ratios. (G) PCA plot of gene expression in AMs stimulated with or without B16 melanoma cells. (H) GO enrichment of gene transcripts upregulated in PBS or IAV AMs upon B16 melanoma cell stimulation as compared with unstimulated cells. (I) Go enrichment of gene transcripts upregulated in B16 tumor cell-stimulated IAV versus PBS AMs. Bar graphs are presented as mean ± SD. Data in **B** and **F** are representatives of three independent experiments with number of mice per group as indicated (n = 3 or 4 mice per group in **B** and **F**). Data in **C**-**E** and **G**-**I** are from one experiment with n = 2 or 3 mice per group as indicated in **C** and **G**. Two-tailed Student t test was performed for comparisons between two groups.

To characterize the responsiveness of IAV-trained AMs against tumor cells, PBS- or IAV-trained AMs were cocultured with B16 melanoma cells. Consisted with the findings in our previous study^12^, we observed more evident B16 tumor cell killing by IAV- than PBS-AMs (Figures S1D and S1E), suggesting enhanced tumor cytotoxic functions in IAV-trained AM. Notably, there were higher levels of pro-inflammatory cytokines, including TNF, IL-12, and IFN-γ, in the supernatants of tumor-stimulated IAV-AMs than PBS-AMs (Figure 1F). These data suggest that IAV-trained AMs have enhanced responsiveness against tumor cells, as compared to naïve AMs from uninfected mice. To determine the transcriptal response of trained AMs to tumor cells, PBS- or IAV-AMs were cocultured with B16 melanoma cells for 48 hours and AMs were then purified for RNA-seq analysis. Transcriptional profiles were drastically changed in both PBS- and IAV-AMs in response to B16 melanoma cell stimulation (Figures 1G, S1F, and S1G), although GO enrichment analysis of gene clusters showed that there were more immune activation-associated gene transcripts increased in IAV-than PBS-AMs in response to tumor cells, including genes related to type II interferon production, tumor necrosis factor production, phagocytosis, and cell killing (Figures 1H and 1I). These data collectively suggest that IAV-trained AMs are hyper-responsive to both bacteria and tumor cells than naïve AMs.

### IAV-trained AMs bear long-lasting transcriptional and epigenetic imprints of IFN-I signaling

IFN-I represents a characteristic cytokine produced early after respiratory viral infections^25^. Indeed, we previously detected acute and abundant IFN-I production in lung tissues of IAV-infected mice^12^. Given the ubiquitous distribution of IFN-I receptor on various cell types including AMs^25^, we speculate that IAV-induced IFN-I stimulates AMs early after infection and thus might contribute to IAV-induced TI in AMs via shaping long-term transcriptional and epigenetic profiles in AMs. To determine whether AMs carry imprints of acute IFN-I stimulation long after IAV infection, we performed RNA-seq analysis in purified AMs at 30 days post IAV infection. We observed that gene transcripts related to response to IFN-I was higher in IAV- than PBS-AMs (Figure 2A). Moreover, assay for transposase-accessible chromatin with high throughput sequencing (ATAC-seq) analysis showed that IAV-AMs had increased chromatin accessibility of 1725 genes and that of 328 genes reduced, as compared to PBS-AMs (Figure 2B). Notably, chromatin accessibility of genes associated with type I interferon signaling pathway was increased in IAV-versus PBS-AMs, including representative interferon-stimulated genes (ISGs) such as *Ifi47*, *Ifitm2*, *Irf7*, *Ifi204* and *Gm4951* (Figures 2C, 2D, and 2E). In the 513 genes with both increased transcription and increased chromatin accessibility in IAV- over PBS-AMs, multiple immune activation-related pathways were enriched, including but not limited to type I and type II IFN responses (Figures 2F and 2G). These data suggest that IAV-induced acute IFN-I signaling leaves epigenetic and transcriptional imprints in AMs long after resolution of IAV-induced respiratory inflammation, which presumably shape TI functions in IAV-trained AMs.

**Figure 2.**
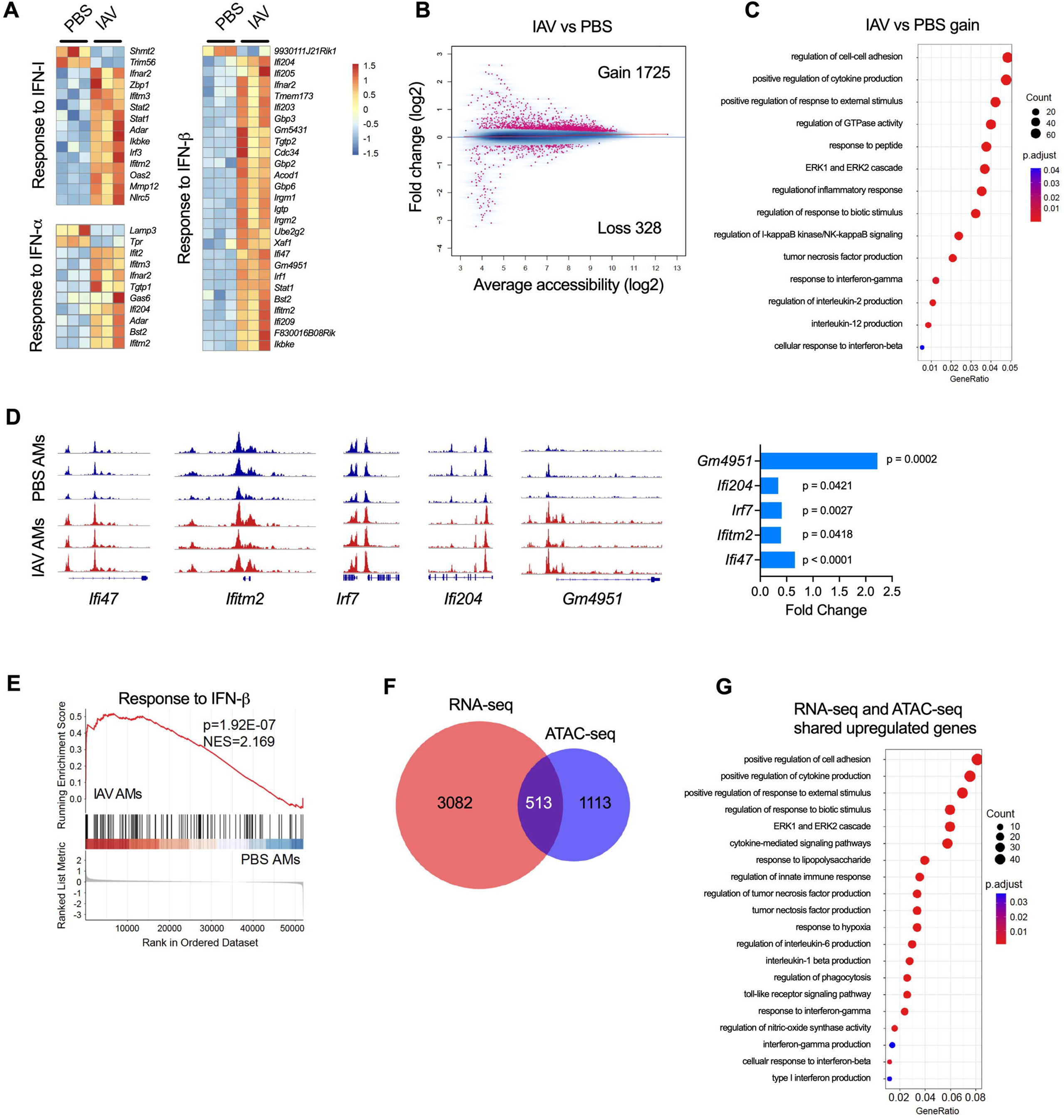
IAV-trained AMs bear long-lasting transcriptional and epigenetic imprints of IFN-I signaling. (A) RNA-seq heatmaps showing z scores of different gene transcripts related to response to IFN-I in AMs from uninfected (PBS AMs) or d30 IAV-infected (IAV AMs) mice. (B) Mean difference versus average (MA) plot showing genes with significant gain or loss of chromatin accessibility in PBS or IAV AMs. (C) GO enrichment of gene clusters with gain of chromatin accessibility in IAV versus PBS AMs. (D) ATAC-seq signals with fold change for genes, including *Ifi47*, *Ifitm2*, *Irf7*, *Ifi204* and *GM4951*, in PBS and IAV AMs. (E) GSEA of ATAC-seq signals of genes related to response to IFN-β in PBS and IAV AMs. (F) Venn plot showing genes with increased transcription (RNA-seq) and/or gain of chromatin accessibility (ATAC-seq) in IAV versus PBS AMs. (G) GO enrichment of clusters of genes with increased transcription (RNA-seq) and gain of chromatin accessibility (ATAC-seq) in IAV versus PBS AMs. Data are from one experiment with n = 3 mice per group.

### IFN-I signaling limits anti-bacterial TI functions in IAV-trained AMs

To determine the impact of IFN-I signaling on the functionality of IAV-trained AMs, wild type (WT) or IFN-I receptor-deficient (IFNAR^-/-^) mice were infected intranasally (i.n.) with IAV, followed by phenotypic and functional analysis on AMs at 30 days post infection (Figures 3A and S2A). IAV infection induces respiratory inflammation, as evidenced by transient body weight loss and production of interferons in lung tissues which was resolved by around 2 weeks after infection in both strains of mice (Figures S3A-S3C). Flow cytometry analysis showed that IAV-exposed AMs in both WT and IFNAR^-/-^mice expressed higher levels of major histocompatibility complex class II (MHC II) than those in uninfected mice, although IAV-exposed IFNAR^-/-^ AMs expressed higher MHC II than IAV-exposed WT AMs (Figure 3B). In response to *ex vivo* LPS or S.P. stimulation, IFNAR^-/-^ IAV-AMs secreted higher amount of pro-inflammatory cyto-/chemokines than WT IAV-AMs (Figure 3C). IFNAR^-/-^ IAV-AMs also showed higher *ex vivo* S.P. bacterial phagocytosis and killing than WT IAV-AMs (Figures 3D and 3E). Notably, IAV-experienced IFNAR^-/-^ mice showed increased resistance to respiratory challenge with S.P. than their WT counterparts, as demonstrated by less body weight loss and lower lung bacterial burdens in IAV-experienced IFNAR^-/-^ than WT mice (Figures 3F and 3G). These data suggest that IFN-I signaling limits IAV-induced anti-bacterial TI functions in AMs. Our previous study suggests that IAV-induced IFN-γ plays critical roles in TI development in AMs^11,12^. To test whether the enhanced anti-bacterial TI functions observed in IFNAR^-/-^ AMs are due to altered IFN-γ signaling in IFNAR^-/-^ mice, we determined the kinetics of both IFN-γ production in lung tissues and expression of CD119 (α chain of IFN-γ receptor) on AMs after IAV infection. We observed comparable concentrations of IFN-γ in lung tissues and median fluorescence intensity (MFI) of CD119 on AMs in WT and IFNAR^-/-^ mice post infection (Figures S3C and S3D). Moreover, we performed intracellular flow cytometry staining of phosphorylated signal transducer and activator of transcription-1 (p-STAT1) and observed comparable levels of intracellular p-STAT1 in AMs from WT and IFNAR^-/-^ mice at 6 days post IAV infection, when both IFN-I and IFN-γ are present abundantly in lung tissues (Figures S3B, S3C, and S3E). These data suggest that altered TI functions in IFNAR^-/-^ AMs are not due to augmented IFN-γ signaling.

**Figure 3.**
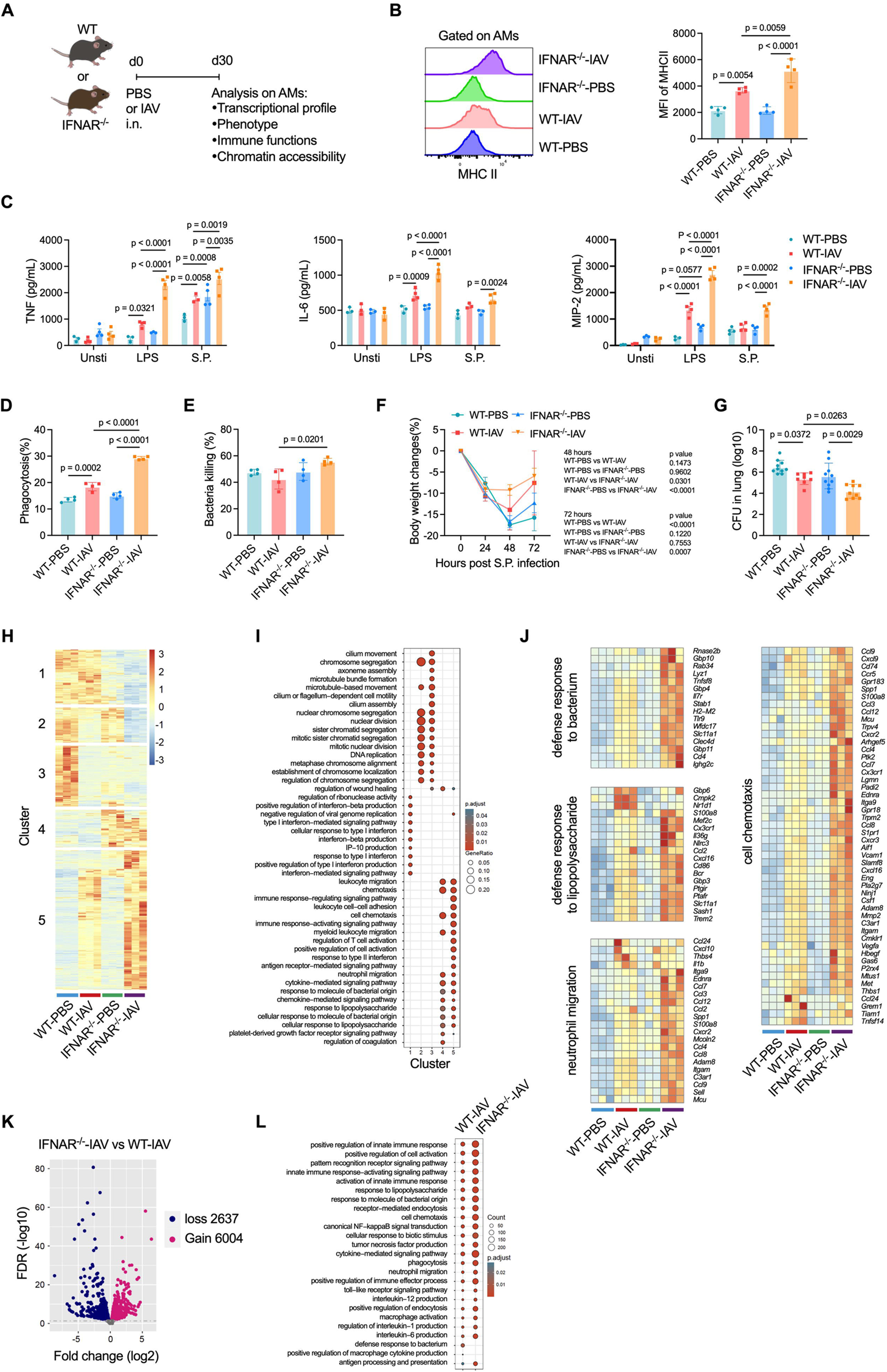
IFN-I signaling limits anti-bacterial TI functions in IAV-trained AMs. (A) Schema of IAV infection in wild type (WT) or type I interferon receptor deficient (IFNAR^-/-^) mice and characterization of AMs on day 30 after infection. (B) Representative flow cytometry histograms and median fluorescence intensity (MFI) of MHC class II expression in PBS and IAV AMs. (C) Concentrations of representative proinflammatory cyto-/chemokines, including TNF, IL-6 and MIP-2, in supernatants of *ex vivo* cultured AMs that are either unstimulated (Unsti) or stimulated with LPS or live *Streptococcal pneumoniae* (S.P.). (D,E) *Ex vivo* bacterial phagocytosis (D) and killing (E) by AMs isolated from WT or IFNAR^-/-^ mice with or without prior IAV infection. (F,G) Percentages of body weight change at 24, 48 and 72 hours post respiratory S.P. infection (F) and bacterial colony forming units (CFU) in lung homogenates at 48 hours post infection (G) in WT and IFNAR^-/-^ mice with or without prior IAV infection. (H) RNA-seq heatmap showing z scores of gene transcripts in AMs from PBS- or IAV- infected mice. (I) GO enrichment analysis of differentially expressed genes shown in **H**. (J) RNA-seq heatmaps showing differentially expressed genes related to selected immune pathways related to anti-bacterial immunity. (K) ATAC-seq volcano plot showing genes with gain or loss of chromatin accessibility in AMs from IAV-infected IFNAR^-/-^ versus WT mice. (L) GO enrichment of genes with gain of chromatin accessibility in AMs from IAV-infected WT or IFNAR^-/-^ mice. Graphs with error bars are presented as mean ± SD. Data in **B** and **C** are representatives of three independent experiments with number of mice per group as indicated (n = 3 or 4 mice per group in **B** and **C**). Data in **D**-**F** are representatives of two independent experiments with n = 4 or 5 mice per group. Data in **G** are pooled from two experiments. Data in **H**-**L** are from one experiment with n = 3 mice per group. One-way ANOVA followed by a Tukey test was performed to compare more than two groups.

We also performed RNA-seq analysis on WT versus IFNAR^-/-^ AMs isolated from uninfected (PBS) or day 30 IAV-infected mice. We found that IFNAR^-/-^-IAV AMs expressed the highest levels of genes related to immune activation, in particular those on anti-bacterial responses including defense response to bacterium and lipopolysaccharide, neutrophil migration, and cell chemotaxis (Figures 3H-3J, S3F, and S3G). To determine the effect of IFN-I signaling on the epigenetic landscape in trained AMs, we performed ATAC-seq analysis in AMs and found increased chromatin accessibility in 6,004 genes and decreased in 2,637 genes in IAV-trained IFNAR^-/-^ versus WT AMs (Figure 3K). GO enrichment of specific marker genes further showed increased chromatin accessibility of genes related to anti-bacterial functions in trained IFNAR^-/-^ than WT AMs (Figure 3L). To test if IFN-I signaling also causes AMs’ metabolic rewiring which is relevant to TI functions^1,4^, we performed *ex vivo* seahorse analysis on oxygen consumption rate (OCR) and extracellular acidification rate (ECAR) in AMs. We found nearly unaltered OCR and ECAR parameters in IAV-trained IFNAR^-/-^ versus WT AMs, indicating a negligible role of IFN-I in the metabolic rewiring in IAV-trained AMs (Figures S3H-S3K). These findings collectively suggest that IFN-I signaling limits IAV-induced anti-bacterial TI functions in AMs, which is associated with IFN-I-dependent epigenetic and transcriptional changes of genes related to anti-bacterial innate immunity.

### IAV-trained AMs in IFNAR^-/-^ mice develop independently of circulating monocytes

Previous studies suggest that ontogeny of AMs, namely embryonic-derived resident AMs (TRAMs) versus monocyte-derived AMs (MoAMs) is a deciding factor on their functionality^20,28,29^. We show in our previous study that IAV-trained AMs in WT mice develop independently of circulating monocytes^12^. It is therefore relevant to determine whether AMs in IAV-experienced IFNAR^-/-^ mice derive from TRAMs or MoAMs, so as to exclude the possibility that the functional differences in WT versus IFNAR^-/-^ trained AMs were due to more evident reconstitution of MoAMs in IFNAR^-/-^ than WT mice. To do so, we utilized a multitude of approaches including phenotypic, transcriptional, and epigenetic analysis on AMs, as well as AM lineage-tracing models. Flow cytometry analysis showed comparable levels of cell surface markers, including CD11b, Ly6C, CD11c, CD64, and Siglec-F, between PBS- and IAV-trained AMs in IFNAR^-/-^ and WT mice, and the expression levels of these markers were drastically different from those of circulating monocytes (Figure 4A). Moreover, RNA-seq analysis in purified AMs showed that gene transcripts of characteristic core transcription factors of AMs, embryonic-derived signature genes, and monocyte-derived signature genes remain unchanged in WT versus IFNAR^-/-^ AMs before or after IAV infection (Figure 4B). ATAC-seq analysis in AMs showed that chromatin accessibility in signature genes of TRAMs, including *Marco*, *Car4*, and *Rxra*^30,31^, was unaltered before and after IAV infection in WT and IFNAR^-/-^ mice (Figure 4C). These data suggest that IAV-trained AMs from both WT and IFNAR^-/-^ mice derived from TRAMs instead of MoAMs.

**Figure 4.**
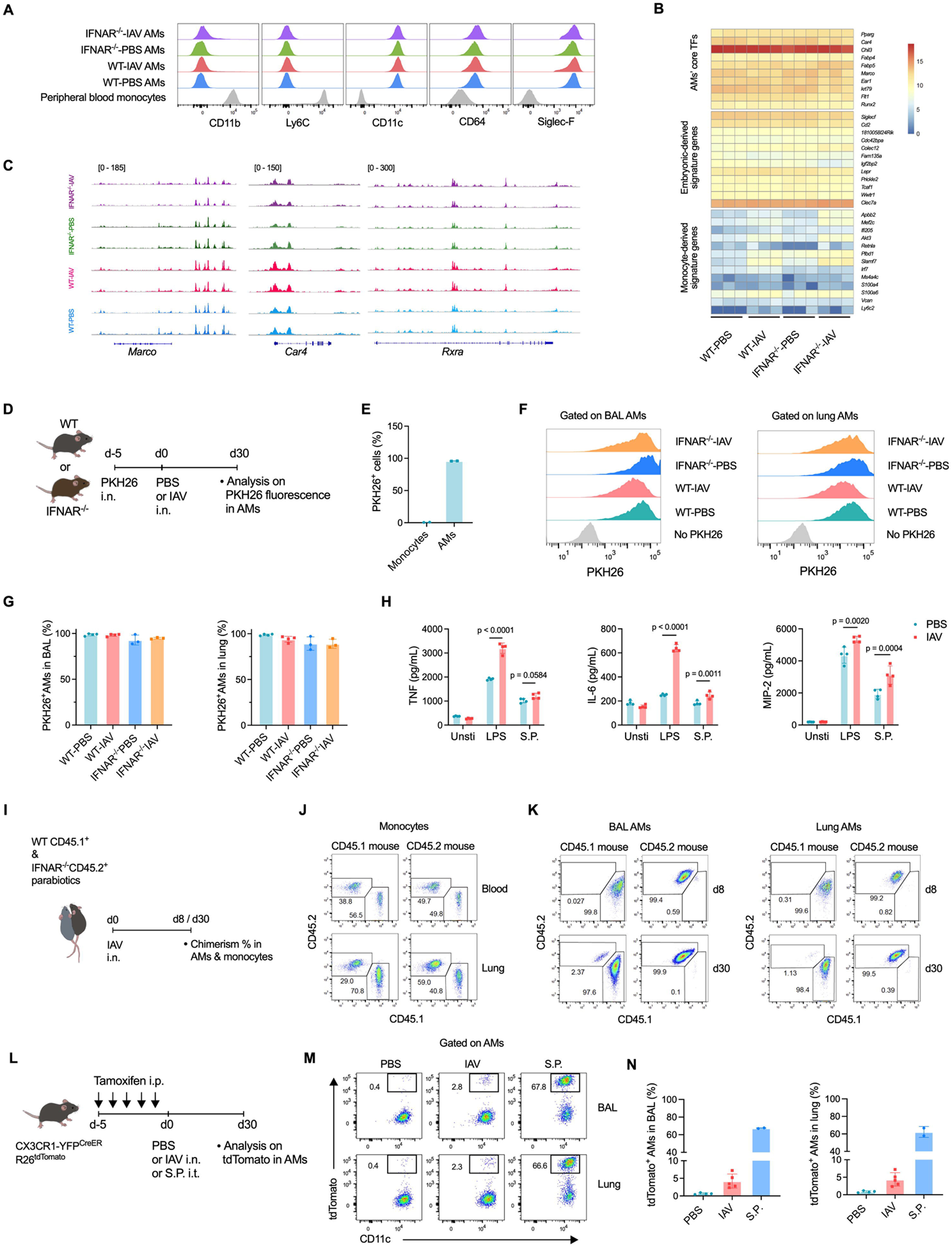
IAV-trained AMs in IFNAR^-/-^ mice develop independently of circulating monocytes. (A) Representative flow cytometry histograms of surface molecules on peripheral blood monocytes and AMs from uninfected (PBS) or IAV-infected WT or IFNAR^-/-^ mice. (B) Heatmaps of RNA-seq data showing genes transcripts of core transcriptional factors (TFs) of AMs, embryonic-derived and monocyte-derived signature genes in AMs. (C) ATAC-seq peaks of *Marco*, *Car4*, and *Rxra* in AMs. (D) Schema of PKH26 staining of AMs before IAV infection in WT or IFNAR^-/-^ mice. (E) Percentage of PKH26^+^ cells in peripheral blood monocytes and AMs immediately before IAV infection. (F) Representative flow cytometry histograms of PKH26 fluorescence in AMs from BAL and digested lung tissues. (G) Percentages of PKH26^+^ AMs in BAL and digested lung tissues. (H) Concentrations of representative proinflammatory cyto-/chemokines, including TNF, IL-6, and MIP-2, in supernatants of *ex vivo* stimulated PKH26^+^ AMs purified from IAV-infected or uninfected (PBS) WT mice. (I) Schema of parabiotic mouse model. (J,K) Representative flow cytometry dot plots showing the chimerisms of circulating monocytes (J) and AMs isolated from BAL and digested lung tissues (K) in parabiotic mice as shown in **I**. (L) Schema of IAV or S.P. infection in an AM fate-mapping Cx3cr1-YFP^CreER^R26^tdTomato^ mouse model. (M,N) Representative flow cytometry dot plots showing tdTomato fluorescence in AMs (M) and percentages of tdTomato^+^ AMs (N) from uninfected (PBS), IAV-infected, or S.P. infected mice at 30 days post infection as shown in **L**. Bar graphs are presented as mean ± SD. Data in **A**, **D**-**K** are representatives of three independent experiments with n = 2, 3, or 4 mice or duplicated culture wells per group. Data in **M** and **N** are representatives of two independent experiments with n = 2, 4, or 5 mice per group. Data in **B** and **C** are from one experiment with n = 3 mice per group. Numbers in **J**, **K**, and **M** indicate percentage against parent gate. Two-tailed Student t test was performed for comparisons between two groups. One-way ANOVA followed by a Tukey test was performed to compare more than two groups.

To further show the cellular origin of AMs post IAV infection, we utilized an AM lineage-tracing model where TRAMs are labelled by i.n. distillation of the PKH26 cell staining dye^16,32^, which preferentially labels AMs but not circulating monocytes (Figures 4D and 4E). At 30 days post IAV infection, over 90% of AMs in both WT and IFNAR^-/-^ mice showed a PKH26^hi^ phenotype (Figures 4F and 4G). These data suggest that the vast majority of post-IAV AMs in both WT and IFNAR^-/-^ mice derived from embryonic-derived TRAMs but not circulating monocytes. We also purified PKH26^hi^ AMs from day 30 IAV-infected or uninfected WT mice, and stimulated them with LPS or live S.P. *ex vivo*. We observed increased proinflammatory cyto-/chemokine production by IAV-experienced PKH26^hi^ TRAMs in response to restimulation (Figure 4H). These data thus support our conclusion that IAV induces TI in embryonic-derived TRAMs. To further show the independence of monocytes in IAV-trained AMs, we infected WT (CD45.1^+^) and IFNAR^-/-^ (CD45.2^+^) parabiotic mice with IAV and examined the levels of chimerism of AMs post infection (Figure 4I). Chimerisms of peripheral blood and monocytes were typically over 30% in both WT and IFNAR^-/-^ parabionts at 30 days post infection (Figure 4J). In sharp contrast, minimal chimerism (∼1% on average) of AMs was observed in either IFNAR^-/-^ or WT mice on days 8 and 30 post infection (Figure 4K), indicating that IAV-trained WT and IFNAR^-/-^ AMs develop independently of circulating monocytes. These data thus suggest that the functional differences in IAV-trained WT versus IFNAR^-/-^ AMs are not due to recruitment of MoAMs.

We have also utilized an AM fate-mapping Cx3cr1-YFP^CreER^R26^tdTomato^ mouse model to determine the cellular ontogeny of IAV-experienced AMs (Figure 4L)^33^. Repeated doses of Tamoxifen in Cx3cr1-YFP^CreER^R26^tdTomato^ mice induced tdTomato expression in nearly 100% of peripheral blood monocytes (Figures S4A-S4C). By 30 days after infection, around 30% of peripheral blood monocytes still showed tdTomato expression (Figures S4B and S4C). Notably, there were minimal (less than 1%) tdTomato^+^ AMs in uninfected mice, which is consistent with data in previous study^33^. In mice infected with IAV, we observed less than 5% of tdTomato^+^ AMs at 30 days post infection, suggesting that the majority of IAV-experienced AMs derive from embryonic-derived TRAMs (Figures 4M and 4N). We recently showed that S.P. infection induces development of MoAMs^34^. We therefore infected Cx3cr1-YFP^CreER^R26^tdTomato^ mice with S.P., and we observed that over 60% of AMs are tdTomato^+^ at 30 days post S.P. infection. These data collectively suggest that IAV infection induces trained immunity in TRAMs independently of MoAMs.

### IFN-I signaling is required to developing anti-tumor TI functions in AMs

We next sought to determine the roles of IFN-I signaling in anti-tumor TI functions. To address this, IAV-infected WT and IFNAR^-/-^ mice at day 30 post infection were injected intravenously (i.v.) with luciferase-expressing B16 melanoma cells (B16-luc) to inoculate tumor cells in the lungs, followed by analysis of lung tumor burdens at endpoint (Figure 5A). Consistent with the findings in our previous study^12^, IAV-infected WT mice had reduced lung tumor burdens as compared to uninfected (PBS) WT mice (Figures 5B-5G). Lung B16 tumor burdens in IFNAR^-/-^ mice were significantly higher than those in WT mice, likely due to impaired NK cell-mediated anti-tumor functions^35^. Importantly, lung tumor burdens were comparable in IAV-infected and uninfected IFNAR^-/-^ mice (Figures 5B-5G). Given the critical roles of IAV-trained AMs in WT mice to exert anti-tumor TI functions (Figures S1D and S1E)^12^, these observations in IFNAR^-/-^ mice point to an indispensable role of IFN-I signaling in developing anti-tumor TI functions in IAV-trained AMs.

**Figure 5.**
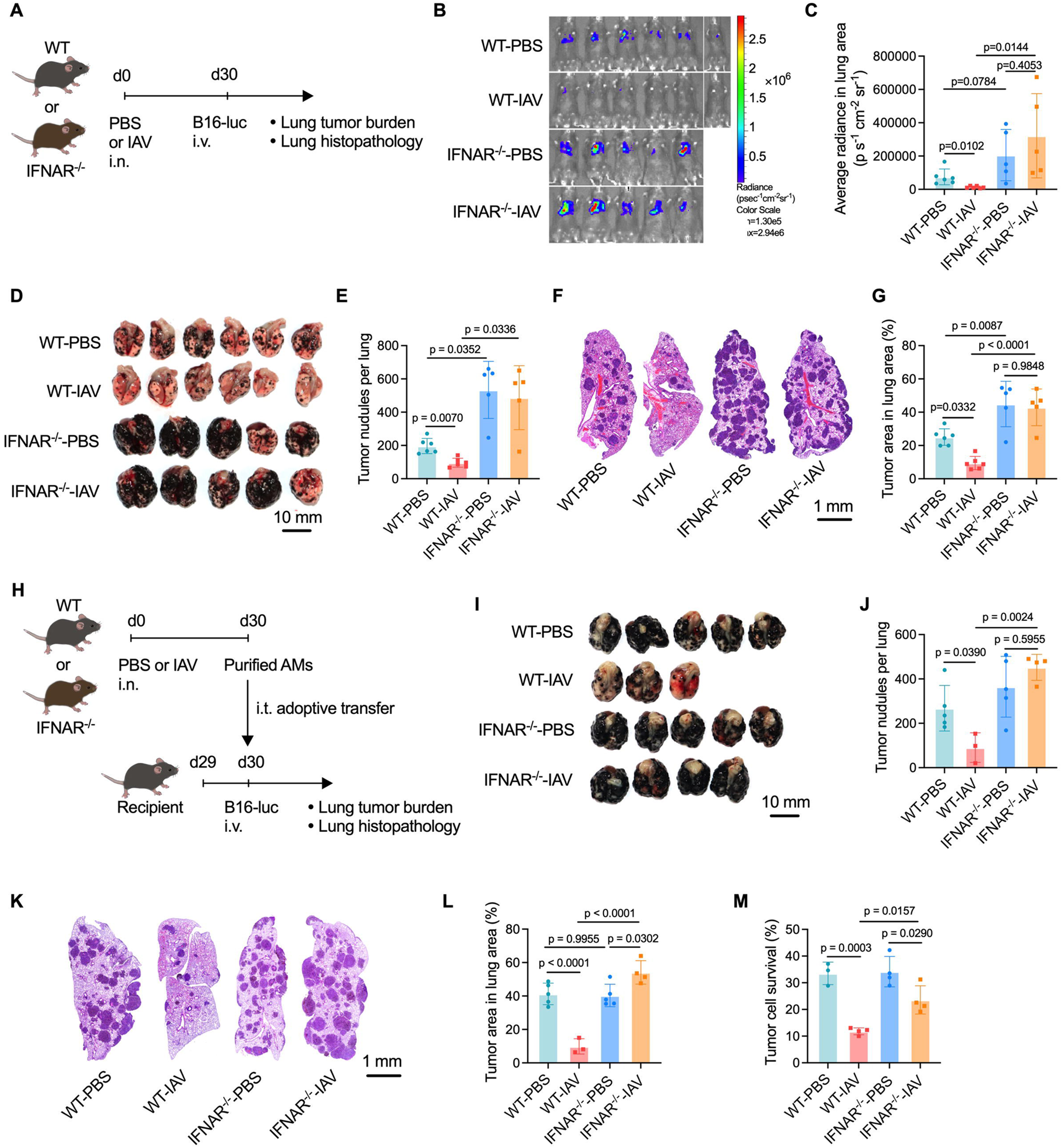
IFN-I signaling is required to developing anti-tumor TI functions in AMs. (A) Schema of IAV infection in WT or IFNAR^-/-^ mice followed by i.v. inoculation of luciferase-expressing B16 melanoma cells (B16-luc). (B, C) Luciferin-based *in vivo* tumor imaging (B) and average radiance of luciferin signals in the lung area in uninfected (PBS) or IAV-infected WT or IFNAR^-/-^ mice (C). (D, E) Macroscopic evaluation of the lungs (D) and number of macroscopically visible B16 tumor nodules on the surface of lung lobes in mice showed in **A** (E). (F, G) Representative lung histopathology images (F) and percentage of lung area occupied by tumor lesions based on lung histopathological analysis in mice shown in **A** (G). (H) Schema of intratracheal (i.t.) adoptive transfer of AMs isolated from PBS- or day 30 IAV-infected WT or IFNAR^-/-^ mice into naive recipient mice, which were i.v. inoculated with B16 melanoma cells to the lungs at one day prior to adoptive transfer of AMs. (I, J) Macroscopic evaluation of the lungs (I) and number of macroscopically visible B16 tumor nodules on the surface of lung lobes of recipient mice shown in **H** (J). (K, L) Representative lung histopathology images (K) and percentage of lung area occupied by tumor lesions in mice shown in **H** (L). (M) Survival of B16 melanoma cells at 72 hour after coculture with PBS or IAV AMs from WT or IFNAR^-/-^ mice at an E:T ratio of 10:1. Bar graphs are presented as mean ± SD. Data are representatives of two independent experiments with number of mice per group as indicated (n = 5 or 6 mice per group in **B**-**G**; n = 3, 4 or 5 mice per group in **I**-**L**; n = 3 or 4 duplicated culture wells per group in **M**). One-way ANOVA followed by a Tukey test was performed to compare more than two groups.

To further show that IFN-I signaling is required to developing anti-tumor TI functions in AMs, we compared the anti-tumor potential of IAV-trained WT versus IFNAR^-/-^ AMs by using both airway adoptive transfer of AMs in lung tumor-bearing mice and *ex vivo* AM-tumor cell coculture for cytotoxicity analysis. To compare *in vivo* anti-tumor functions of IAV-trained WT versus IFNAR^-/-^ AMs in a therapeutic scenario, naïve WT recipient mice were injected intravenously (i.v.) with B16 melanoma cells. One day after tumor inoculation, recipient mice were adoptively transferred intratracheally (i.t.) with AMs isolated from uninfected or day 30 IAV-infected WT or IFNAR^-/-^ mice, followed by analysis of tumor burdens in the lungs (Figure 5H). Lung macroscopic and histopathological analyses at experimental endpoint showed that mice receiving trained WT but not IFNAR^-/-^ AMs had reduced lung tumor burdens (Figures 5I-5L). We also isolated AMs from PBS or day 30 IAV-infected WT or IFNAR^-/-^ mice and cocultured them respectively with B16 melanoma cells, followed by tumor cell survival analysis. Reduced survival was observed in tumor cells coculture with IAV-trained than IAV-naive IFNAR^-/-^ AMs, though trained IFNAR^-/-^ AMs showed reduced tumor cytotoxicity than trained WT AMs (Figure 5M). To determine whether the impact of IFN-I signaling in IAV-trained AMs is exerted during the priming phase (i.e., acute phase after IAV infection) or during the secondary response phase, we administered a single dose of anti-IFNAR1 monoclonal antibody (mAb) into WT mice before IAV infection, and the functionality of IAV-trained AMs were determined at 30 days post infection (Figure S5A). We observed that IAV-trained AMs from mice receiving anti-IFNAR1 mAb showed increased production of proinflammatory cyto-/chemokines upon *ex vivo* stimulation with LPS, as compared to those without anti-IFNAR1 mAb (Figure S5B). We also inoculated B16 melanoma cells i.v. in mice at 30 days post IAV infection with or without single dose of anti-IFNAR1 mAb before infection (Figure S5C). We observed that blockade of IFN-I signaling during the acute phase of IAV infection impaired IAV-induced anti-tumor immunity in the lung (Figures S5D and S5E). Collectively, these findings suggest that that IFN-I priming signal is required to developing anti-tumor TI functions in IAV-trained AMs.

### IFN-I signaling induces anti-tumor TI via enhanced IFN-**γ** production by trained AMs against tumor cells

To further show how IFN-I signaling promotes anti-tumor TI functions in AMs, we performed RNA-seq analysis in B16 tumor cell-stimulated WT or IFNAR^-/-^ AMs from PBS- or day 30 IAV-infected mice. Three-dimensional principal component analysis (PCA) of RNA-seq data revealed that WT-IAV AMs had more evident transcriptional changes in response to tumor cells than IFNAR^-/-^-IAV AMs, with about 80% variance in PC1 in WT-IAV AMs (WT-IAV AMs+B16 versus WT-IAV AMs) and 20% in IFNAR^-/-^-IAV AMs (IFNAR^-/-^-IAV AMs+B16 versus IFNAR^-/-^-IAV AMs) (Figure 6A). The PC2 axis, which distinguished WT- from IFNAR^-/-^-IAV AMs, accounted for only around 15% of total variance (Figure 6A). Indeed, WT-IAV AMs had only 279 gene transcripts upregulated and 398 down compared to IFNAR^-/-^-IAV AMs (Figure 6B). Such transcriptional differences were enlarged after B16 tumor cell stimulation, with 1673 gene transcripts upregulated and 798 down in WT-versus IFNAR^-/-^-IAV AMs + B16 (Figure 6C). Further analysis on the differentially expressed genes showed that B16 tumor cell stimulation induced upregulation of immune activation-related gene transcripts in WT- but not IFNAR^-/-^-IAV AMs, including gene clusters on activation of innate immune response, positive regulation of type II interferon production, response to type II interferon, and positive regulation of cell killing, indicating enhanced transcriptional anti-tumor response by WT- but not IFNAR^-/-^- IAV AMs (Figures 6D and 6E). These results suggest that IFN-I-signaling is required to developing transcriptional anti-tumor response in trained AMs.

**Figure 6.**
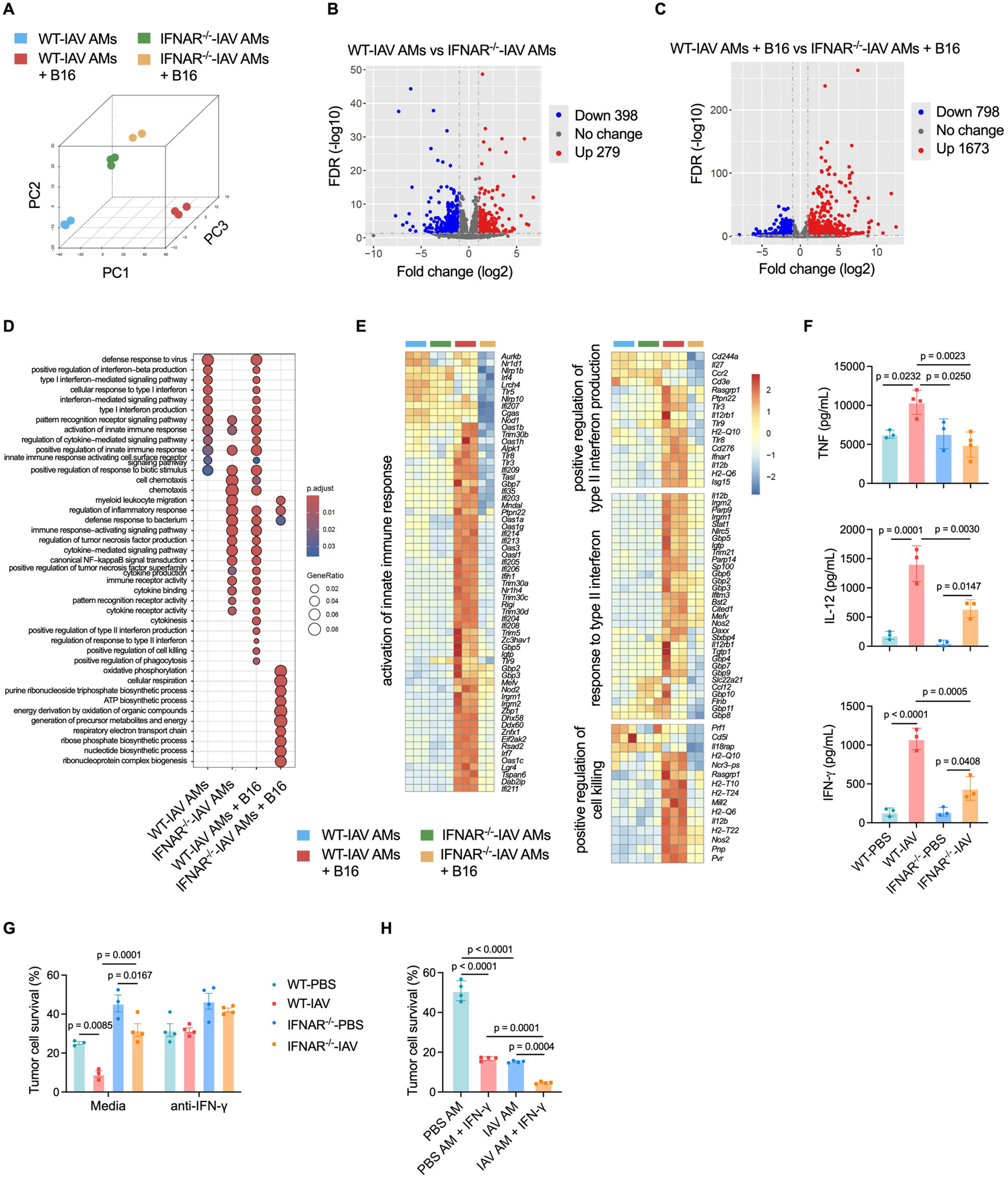
IFN-I signaling induces anti-tumor TI via enhanced IFN-γ production by trained AMs against tumor cells. (A) Three-dimensional PCA plot of RNA-seq data in WT or IFNAR^-/-^ IAV-trained AMs cocultured with or without B16 melanoma cells. (B) Volcano plot showing differentially expressed genes in WT versus IFNAR^-/-^ IAV-trained AMs. (C) Volcano plot showing differentially expressed genes in WT versus IFNAR^-/-^ IAV-trained AMs cocultured with B16 melanoma cells. (D) GO enrichment analysis showing representative functional pathways in WT or IFNAR^-/-^ IAV- trained AM cocultured with or without B16 melanoma cells. (E) RNA-seq heat maps showing differentially expressed genes related to activation of innate immune response, type II interferon production and response, and anti-tumor functions in WT or IFNAR^-/-^ IAV-trained AM cocultured with or without B16 melanoma cells. (F) Concentrations of representative proinflammatory cytokines, including TNF, IL-12 and IFN-γ, in the supernatants of B16 tumor cell-stimulated PBS- or IAV-trained AMs isolated from WT or IFNAR^-/-^ mice. (G) Survival of B16 melanoma cells cocultured with PBS- or IAV-trained AMs isolated from WT or IFNAR^-/-^ mice, with or without supplementation of anti-IFN-γ blocking antibodies to culture media. (H) Survival of B16 melanoma cells cocultured with PBS- or IAV-trained AMs from WT mice, with or without supplementation of recombinant IFN-γ to culture media. Bar graphs are presented as mean ± SD. Data in **A**-**E** are from one experiment with n = 3 biological replicates per group in groups including WT-IAV AMs, IFNAR^-/-^-IAV AMs, and WT-IAV AMs + B16, and n = 2 biological replicates per group in IFNAR^-/-^-IAV AMs + B16 group. Data are representatives of three independent experiments in **F**-**H** with n = 3 or 4 duplicated culture wells per group as indicated. One-way ANOVA followed by a Tukey test was performed to compare more than two groups.

To screen key molecular mediators accounting for IFN-I-induced anti-tumor TI functions in AMs, we cocultured B16 melanoma cells with WT or IFNAR^-/-^ AMs isolated from IAV-infected or uninfected mice, followed by quantification of selected pro-inflammatory cytokines in the supernatants of cell culture. While comparable concentrations of TNF, IL-12, and IFN-γ were detected in the supernatants of tumor-stimulated WT- and IFNAR^-/-^-PBS AMs, much higher concentrations of these cytokines were detected in the supernatants of WT- than IFNAR^-/-^-IAV AMs (Figure 6F). These data thus suggest that IFN-I-induced anti-tumor TI functions in AMs are associated with enhanced proinflammatory cytokines produced by trained AMs.

Previous study showed IFN-γ as a critical cytokine that can boost macrophage-mediated anti-tumor functions^36^, and our current experimental data showed that tumor cell stimulation induced more evident transcription of genes related to secretion of and response to IFN-γ, as well as enhanced IFN-γ protein contents in WT- than IFNAR^-/-^-IAV AMs (Figures 6D-6F, S6A, and S6B). Moreover, there were increased contents of IFN-γ but not TNF, IL-12, or IL-6 in the BAL fluid of mice at 16 days after i.v. inoculation of B16 melanoma cells (Figure S7A). In contrast, there were reduced content of IFN-γ in BAL fluid of 48-hour S.P.-infected WT mice with or without prior IAV infection (Figure S7B). These data support an idea that trained AMs preferentially secrete IFN-γ against tumor cells. We therefore speculate that IFN-I-induced anti-tumor TI functions in AMs is related to their enhanced potential of IFN-γ secretion in response to tumor cells. To test this hypothesis, we cocultured WT- or IFNAR^-/-^-IAV AMs with B16 melanoma cells, with IFN-γ-blocking monoclonal antibodies or recombinant IFN-γ supplemented to selected culture wells, to determine tumor cytotoxicity by AMs. We observed that blocking of IFN-γ abrogated the enhance tumor cytotoxicity by trained AMs (Figure 6G), and recombinant IFN-γ boosted tumor cytotoxic effects by both PBS- and IAV-AMs (Figure 6H). Collectively, these results suggest that IFN-I signaling induces anti-tumor TI functions in AMs which produce enhanced IFN-γ against tumor cells.

### IFN-I limits anti-bacterial and induces anti-tumor TI functions in *ex vivo* trained AMs

To determine whether and how *ex vivo* IFN-I priming signal shapes TI functions in AMs, we cultured freshly isolated AMs from naïve WT mice and trained them *ex vivo* with IFN-β in the presence or absence of IFN-γ, a key cytokine required to induce TI in AMs^11,12^. AMs were allowed to rest for 6 days after stimulation with IFNs^4,37^, before re-stimulation with either live S.P./LPS or B16 melanoma cells (Figure 7A). Compared with AMs trained with IFN-β + IFN-γ, those trained with IFN-γ alone secreted higher amount of pro-inflammatory cyto-/chemokines, including TNF, IL-6, and MIP-2, but not IFN-γ, in response to live S.P. or LPS (Figure 7B). In contrast, AMs trained with IFN-γ + IFN-β showed enhanced tumor cytotoxic effects, as compared to those trained with IFN-γ alone (Figure 7C). AMs trained with IFN-β alone did not produce increased proinflammatory cytokines in response to re-stimulation with LPS or live S.P., though they exert enhanced tumor cytotoxic functions as compared to untrained and IFN-γ-trained AMs (Figures 7B and 7C). These data from *ex vivo* cytokine-mediated training system further demonstrate that IFN-I priming signal limits anti-bacterial TI functions but is required to developing anti-tumor TI functions in AMs.

**Figure 7.**
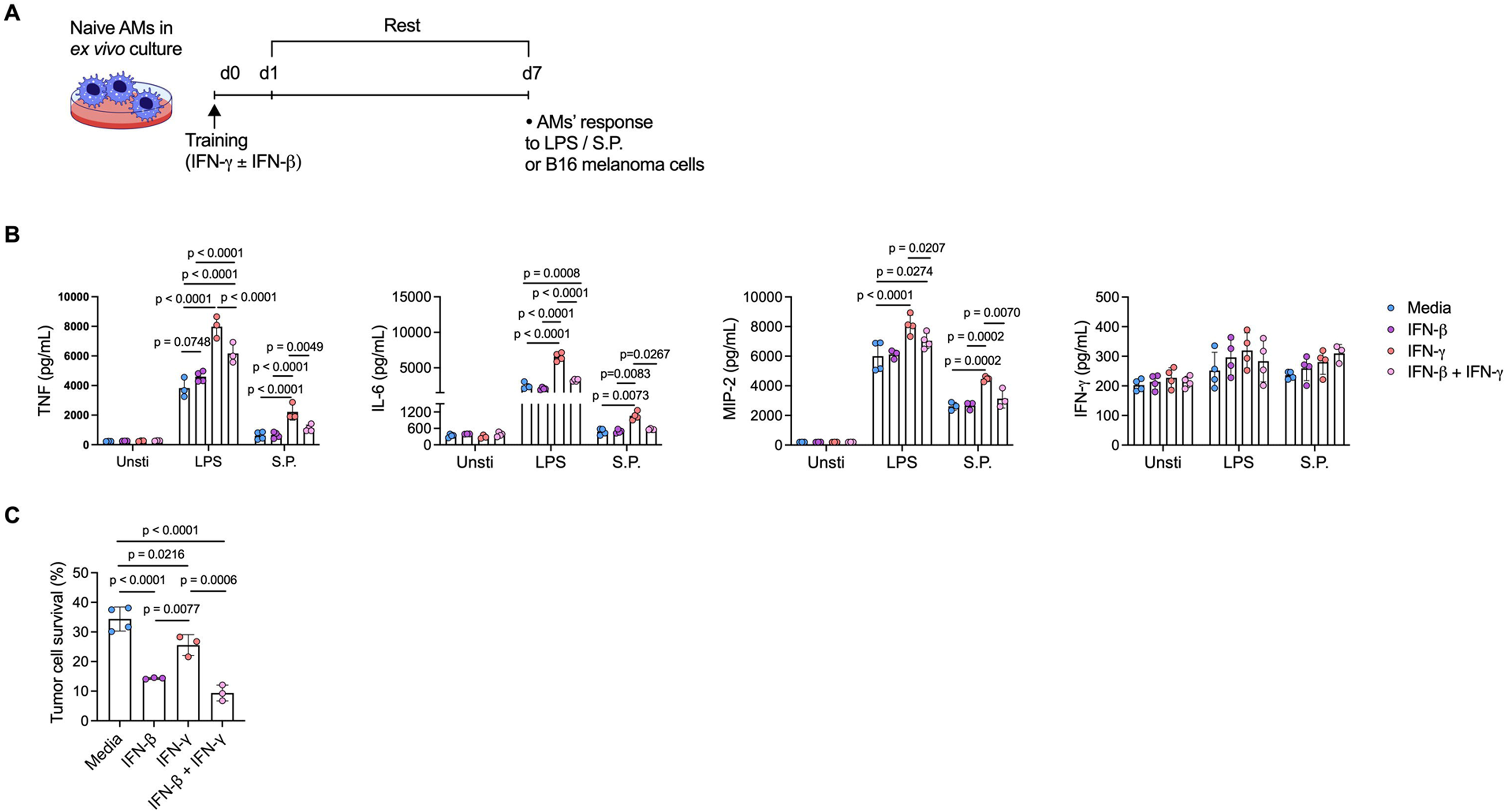
IFN-I limits anti-bacterial and induces anti-tumor TI functions in *ex vivo* trained AMs. (A) Schema of ex vivo training of AMs isolated from wild type naïve mice. Freshly isolated AMs were stimulated with IFN-γ and/or IFN-β for 24 hours, followed by a rest for 6 days before re- stimulation with LPS, live *Streptococcal pneumoniae* (S.P.), or B16 melanoma cells. (B) Concentrations of representative proinflammatory cytokines, including TNF, IL-6, MIP-2 and IFN-γ, in supernatants of *ex vivo* trained AMs stimulated with LPS or S.P.. (C) Survival of B16 melanoma cells after coculture with *ex vivo* trained AMs. Bar graphs are presented as mean ± SD. Data are representatives of two independent experiments with n = 3 or 4 duplicated culture wells per group as indicated. One-way ANOVA followed by a Tukey test was performed to compare more than two groups.

### Human AMs carrying transcriptional imprints of IFN-I signaling are associated with enhanced anti-tumor immunity

Our data thus far suggest that acute IFN-I signaling in mice induces long-lasting epigenetic and transcriptional imprints of prior IFN-I stimulation in AMs which are related to anti-tumor TI functions. We next sought to assess whether trained AMs with IFN-I imprints also exist in human lung tissues and are associated with anti-tumor functions. By using published original data of single-cell RNA sequencing analysis (scRNA-seq) in bronchoalveolar lavage samples from ten healthy human donors^38^, we identified thirteen populations (clusters 0-12) of leukocytes including nine subpopulations of AMs (clusters 0, 1, 2, 4, 5, 6, 7, 8 and 9, designated respectively as m1-m9) based on coexpression of genes characteristic to human AMs including *Mrc1*, *Pparg*, *Serpina1* and *Marco* (Figures 8A and S8A-S8C). Notably, m1 and m5 subpopulations demonstrated transcriptional profiles reminiscent of TI in mouse resident AMs (Figure 8B). GSEA enrichment analysis showed that, compared to m1, m5 had higher gene transcripts related to type I interferon signaling characterized by increased transcripts of ISGs (Figures 8C and 8D). Moreover, GO enrichment of cluster specific marker genes showed enrichment of anti-tumor immunity-related genes in m5 versus m1 populations, including gene clusters related to cell killing, phagocytosis, tumor necrosis factor production and nitric oxide metabolic process (Figure 8E). Similar to our observations in mouse AMs, the increased transcription of IFN-I signaling-related genes in m5 than m1 AMs are associated with increased transcription of genes related to response to IFN-γ, cell killing, and phagocytosis (Figure 8F). Moreover, by using published original scRNA-seq data in non-small cell lung cancer (NSCLC) tissues^39^, we further showed that transcriptional IFN-I imprints in AMs infiltrating NSCLC tissues are associated with anti-tumor immune activation in tumor microenvironment (Figures 8G and 8H). Collectively, these results suggest that trained human AMs carrying imprints of prior IFN-I-signaling have enhanced anti-tumor TI functions at transcription level and are associated with anti-tumor immunity in human lung cancer tissues.

**Figure 8.**
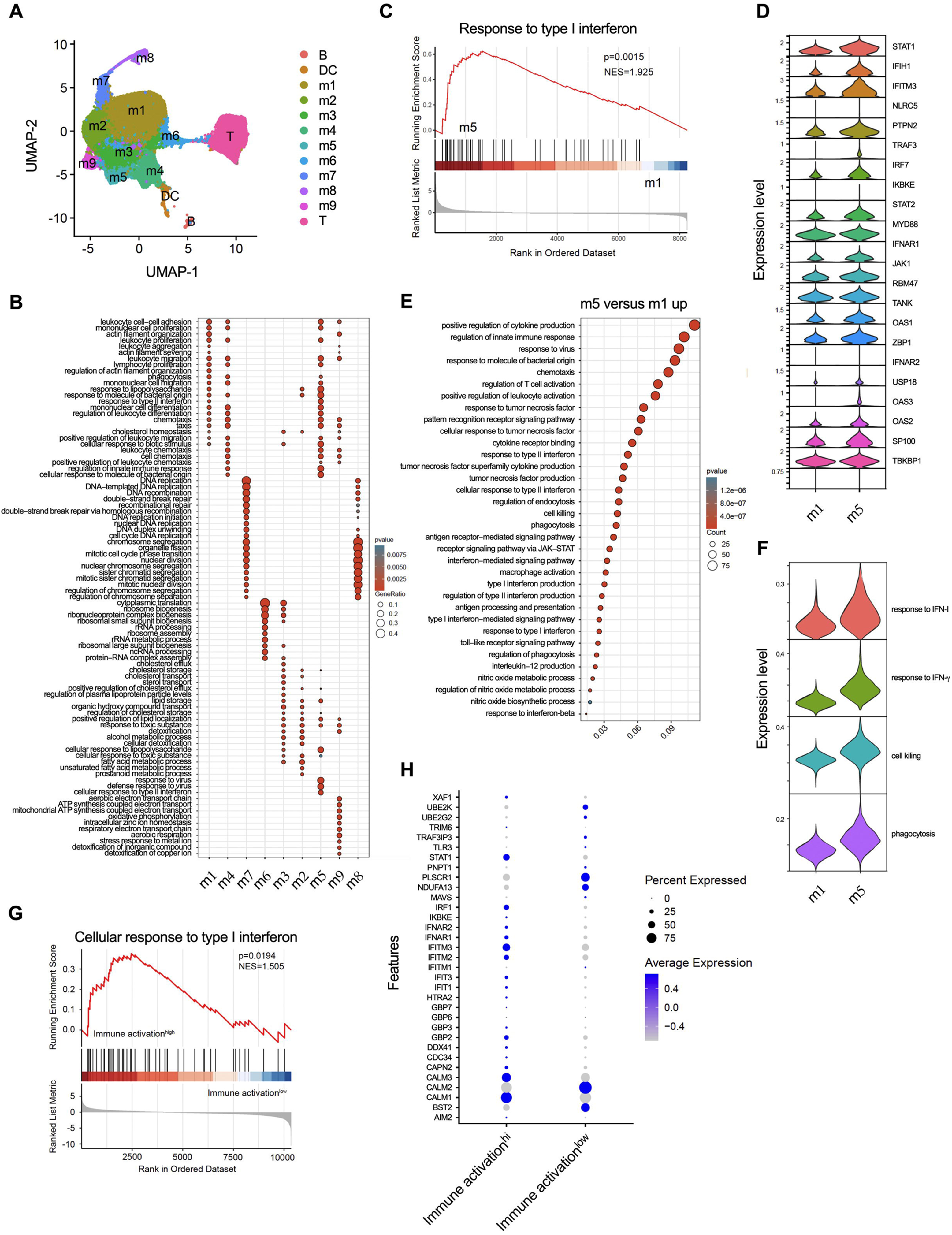
Human AMs carrying transcriptional imprints of IFN-I signaling are associated with enhanced anti-tumor immunity. (A) scRNA-seq UMAP plot showing 13 leukocyte populations in BAL fluid of healthy human donors, including B cells, T cells, dendritic cells (DC), and a total of 9 subpopulations of AMs (m1-m9). (B) GO enrichment of functional gene clusters in subpopulations of AMs. (C) GSEA enrichment of genes related to response to IFN-I in m5 versus m1 subpopulations of AMs. (D) Expression levels of differentially expressed ISGs genes with higher transcription levels in m5 than m1 AMs. (E) GO enrichment of immune functional pathways with gene transcripts upregulated in m5 versus m1 AMs. (F) Expression levels of differentially expressed gene clusters related to response to IFNs and tumor cytotoxic functions in m1 and m5 AMs. (G) GSEA enrichment of scRNA-seq data showing cellular response to IFN-I in trained human AMs from non-small cell lung cancer (NSCLC) tissues with immune activation high or low microenvironment. (H) Transcription levels of genes related to response to IFN-β in NSCLC tissues shown in **G**.

## Discussion

Numerous clinical and experimental studies have linked TI to enhanced host defense against both infections and tumors^6–10,13–16,40–47^. Clinical studies suggest that BCG-induced TI are associated with protection against antigenically-irrelevant pathogens and regression of bladder cancer^8,13,14,46^. TI induced by β-glucan and respiratory viruses was also shown to exert anti-bacterial and anti-tumor responses in experimental animals^10–12,47^. Despite these intriguing and important findings highlighting the roles of TI in both anti-infectious and anti-tumor immunity, much less is known about the key priming signals that respectively contribute to anti-infectious and anti-tumor TI functions. Here, we show that IFN-I signaling during the acute phase of respiratory IAV infection is required to anti-tumor TI development whereas it limits anti-bacterial TI functions in lung tissue-resident AMs. Our study reveals a previously unappreciated mechanism through which anti-tumor TI functions in AMs are preferentially induced by IFN-I priming signal.

Beyond its critical role in anti-viral immunity, IFN-I is a key immune modulator in a wider spectrum of immune scenarios including anti-bacterial immunity, anti-tumor immunity and autoimmunity^25,48–50^. Specifically, IFN-I was shown to acutely promote anti-tumor functions of various immune cells including macrophages, dendritic cells, NK cells, and T cells^49^. Moreover, IFN-I and IFN-I-inducing agents has been tested clinically as potential therapeutics against various human cancers^48,51^. Although these important findings shed light on the significance of IFN-I in anti-tumor immunity, the potential roles of IFN-I in modulating long-term anti-tumor innate immunity (i.e., anti-tumor TI functions) remain unclear. To our best knowledge, our findings show for the first time that IFN-I priming signal is required to long-term innate anti-tumor functions in self-sustaining lung tissue-resident AMs.

We show in previous studies that respiratory viral infection-induced acute IFN-γ production by activated CD8^+^ T cells and NK cells is respectively required to the development of anti-bacterial and anti-tumor TI in AMs^11,12^. An intriguing finding in our current study is that while IFN-I is critically required to the development of anti-tumor TI functions in AMs, it limits their anti-bacterial TI functions. These findings on the differential roles of IFN-γ and IFN-I in inducing and shaping TI functions of AMs collectively point to a hypothesis that, while IFN-γ serves as a key priming signal that triggers the development of TI in AMs with anti-bacterial functions, IFN-I represents a key “ingredient” steering IFN-γ-induced TI in AMs toward anti-tumor functional direction but partially away from anti-bacterial direction. Indeed, in our *ex vivo* training system, AMs trained with IFN-γ alone developed TI with anti-bacterial but not anti-tumor functions, and AMs trained with IFN-γ plus IFN-I bear anti-tumor TI functions but showed less evident anti-bacterial TI functions.

Previous studies suggest that influenza-induced IFN-I impairs anti-bacterial functions of AMs in the acute phase (i.e., one week) after IAV infection, via mechanisms not yet fully understood^26,52^. In this regard, IFN-I was shown to induce methyltransferase Setdb2-mediated H2K9me3 chromatin marks in the promoter region of neutrophil-recruiting chemokine genes such as CXCL1, causing reduced production of CXCL1 by macrophages and subsequently impaired neutrophil chemotaxis upon bacterial infection^52^. Moreover, studies show that BCG-induced IFN-γ induces central TI in bone marrow that boost anti-infectious functions of monocytes/macrophages, whereas *Mycobacterium tuberculosis*-induced IFN-I impairs such an anti-infectious central TI induced by IFN-γ^15,18^. These studies collectively suggest a suppressing role of IFN-I in both acute and long-term anti-bacterial functions of macrophages. IFN-I-induced epigenetic changes were shown to be inheritable during cell division, and persistent epigenetic reprogramming represents a key feature and molecular basis for TI^5,27^. We show in our ATAC-seq data that acute IFN-I signaling is related to long-lasting changes in chromatin accessibility of both ISGs and non-ISG genes in IAV-trained AMs. Such an IFN-I-induced lasting epigenetic reprogramming in AMs are associated with transcriptional and functional changes toward anti-tumor but not anti-bacterial trained immunity. It will be important to determine in future studies how IFN-I preferentially primes anti-tumor but not anti-bacterial TI in AMs. Moreover, it will be important to determine whether IFN-β and IFN-α play overlapping or differential roles in regulating TI functions in AMs.

We previously showed that IAV-trained AMs have enhanced tumor phagocytosis and cytotoxicity both *in vivo* and *ex vivo*^12^. Our data in this study suggest that trained AMs mediate IFN-γ response preferentially against tumor cells but not bacterial stimulation. Moreover, blockade of IFN-γ in the *ex vivo* coculture of trained AMs and B16 melanoma cells completely abrogated the enhanced anti-tumor functions by trained AMs. As IFN-γ was shown to boost tumor cell phagocytosis and killing by tumor-associated macrophages^53^, we speculate that the enhanced tumor phagocytosis and cytotoxic functions of trained AMs are dependent on and induced by their increased IFN-γ production in autocrine manner.

Adoptive transfer of AMs via airway route represents a potential adoptive immunotherapy against lung diseases including malignancies^54,55^. However, AMs from SPF mice were previously shown to exert pro-tumoral in the lung^56,57^, and innate training is critically required for AMs to exert improved host defense functions. Induction of anti-tumor TI in AMs therefore bears the potential for adoptive therapy against pulmonary malignancies. We show in this study that *ex vivo* cultured AMs can be trained by stimulating jointly with IFN-β and IFN-γ to develop anti-tumor TI functions. We also show that airway adoptive transfer of IAV-trained AMs in mice serve as a therapeutic anti-tumor strategy in the lung. These findings highlight the therapeutic potential of trained AMs against pulmonary malignancies. Previous studies suggest that AMs can be cultured *ex vivo* for continuous expansion in the presence of GM-CSF, and these *ex vivo* expanded AMs maintain their identity as lung-resident macrophages^58^. It will be intriguing and important in future studies to test the potential of *ex vivo* expanded and cytokine-trained AMs for adoptive immunotherapy against respiratory malignancies.

One of the limitations in our study is the deficiency of IFN-I signaling in all cell types and at both priming and recall response phases of TI in IFNAR-/- mice. We have utilized alternative experimental strategies, including the antibody-mediated blocking of IFNAR1 during the priming phase but not recall response phase of TI in AMs, and an *ex vivo* AM training system with IFN-I and/or IFN-γ. Our data from these experiments further support the roles of IFN-I priming signal in shaping TI functions in AMs. However, further studies utilizing AM-specific IFNAR knockout mice are required to more directly show the roles of IFN-I signal specifically in AMs.

In conclusion, our findings shed light on the critical roles of IFN-I signaling, in addition to other key TI-inducing signals such as IFN-γ, in the development of versatile TI functions in AMs with balanced anti-bacterial and anti-tumor innate immunity. Our study will foster future investigation on IFN-I-dependent training of AMs to be exploited as a potential lung tissue specific anti-tumor strategy.

## Acknowledgements

This work was supported by National Natural Science Foundation of China (31970861, 32170893, 32470947, 32400742, and 32100714), Zhejiang Provincial Natural Science Foundation (LR20H100001 and LRG25H100001), China Postdoctoral Science Foundation (2024M752824 and GZC20241486). We thank Professors Zhou Xing (McMaster University) for proofreading the manuscript and helpful discussion. We thank Professors Xuetao Cao and Zhenhong Guo (National Key Laboratory of Medical Immunology), Professors Jianli Wang Qingqing Wang, Feng Xu, Xiaojian Wang, Zhijian Cai, and Xiao Shen (Zhejiang University) providing experimental materials or technical assistance. We thank Yanwei Li and Yueting Xing from the Core Facilities of Zhejiang University School of Medicine for research instrument services and technical assistance.

## Author contributions

T.W. and Y.Y. conceived and designed the study. T.W., J.Z., Y.M, Y.W., Y.L., L.W., Y.L., and C.C. performed experiments. E.C. provided experimental materials. T.W. and Y.Y. analyzed the data and wrote the paper.

## Methods

### Mice

Wild-type female C57BL/6 mice were purchased from Slac Laboratory Animals (Shanghai, China) and were 6-8 week of age upon arrival. CD45.1 congenic mice on a C57BL/6 background (B6.SJL-Ptprc^a^ Pepc^b^/BoyJ) were originally from the Jackson Laboratory (Bar Harbor, ME) and were a kind gift from Professor Zhijian Cai from Zhejiang University. Type I interferon receptor (IFNAR) knockout mice on a C57BL/6 background (B6.129S2-*Ifnar1^tm1Agt^*/Mmjax) were originally from the Jackson Laboratory (Bar Harbor, ME) and were a kind gift from Professor Xiaojian Wang from Zhejiang University. Cx3cr1-YFP^CreER^ mice (B6.129P2(Cg)-*Cx3cr1*^tm2.1(cre/ERT2)Litt^/WganJ, Jax 021160) and R26^tdTomato^ mice (B6.Cg-Gt*(ROSA)*26Sor^tm14(CAG-tdTomato)Hze^/J, Jax 007914) were originally from the Jackson Laboratory (Bar Harbor, ME) and were kind gifts from Professor Xiao Shen from Zhejiang University. Mice were housed in specific pathogen-free facility at Zhejiang University Laboratory Animal Center. All experiments were carried out in accordance with the guidelines from the Animal Research and Ethics Boards of Zhejiang University.

### Respiratory viral and bacterial infection

Acute respiratory viral infection was introduced by intranasal administration of influenza virus (A/PR8/34; 120 pfu per mouse) as previously described^12^. Mice were anesthetized with isoflurane and infected intranasally with 25 μL of virus preparation in PBS. Growth and infection of mice with *Streptococcus pneumoniae* serotype 3 (ATCC 6303) were conducted as previously described^11^. For bacterial resuscitation, frozen bacterial stocks were revived by plating on blood tryptic soy agar containing 5% defibrinated sheep blood (YC Biotech) and 10 mg/ml neomycin, followed by overnight incubation at 37 in a 5% CO_2_ incubator. Resultant colonies were resuspended and grown in Todd Hewitt broth (BD Biosciences) at 37 with 5% CO_2_. Prior to infection, bacteria were harvested and resuspended in PBS. Quantification of colony-forming units (CFUs) was performed by plating 10-fold serial dilutions of bacteria on blood-agar plates that were cultured overnight at 37 °C in 5% CO_2_. Respiratory infection with *Streptococcus pneumoniae* was introduced by intratracheal administration of bacteria resuspended in 40 μL of PBS at 5 × 10^7^ CFUs per mouse.

### Mouse tumor model

GFP-expressing B16F10 melanoma cells (B16-GFP) were generated from luciferase-expressing B16F10 (B16-luc; ATCC CRL-6475-LUC2) by transduction with lentiviral vector encoding GFP gene under EF-1 alpha promoter. B16F10 (B16; ATCC CRL-6475), B16-luc, and B16-GFP cells were grown in RPMI 1640 cell culture medium (Thermo Fisher, Waltham, MA) supplemented with 10% fetal bovine serum (Serana, Brandenburg, Germany) and 1% penicillin/streptomycin (Biosharp, Hefei, China). A total of 5 × 10^5^ B16-luc cells were resuspended in 200 μL PBS, followed by intravenously injected into the tail vein to establish a murine lung melanoma inoculation model.

### Luciferin-based *in vivo* tumor imaging

Mice were anesthetized with isoflurane and injected intraperitoneally with 150 mg/kg D-luciferin (Yeason Biotechnology, Shanghai, China). After injection, mice were placed on the imaging platform of an IVIS Spectrum imaging station (PerkinElmer, Waltham, MA) with continuous isoflurane inhalation anesthesia at 5% with 1 L/min flow of oxygen during the imaging procedure. White light and luciferase activity were recorded for 40 seconds starting at 8 minutes post D-luciferin injection. Images were subjected to interpretation using Living Image software (PerkinElmer, Waltham, MA) for quantification of luciferase activity.

### Lung macroscopy and histopathology

Lung lobes were harvested at desired time points between days 15 and 18 post tumor inoculation. Visible B16 tumor nodules on the surface of all lung lobes were counted under a JSZ8 dissection microscope (Jiangnan Novel Optics, Nanjing, China). Lung lobes were inflated with 10% formalin for 72 hours, followed by embedding in paraffin and subsequent H&E staining for histological examination. Stained lung sections were scanned on a VS200 research slide scanner (Olympus, Tokyo, Japan) by using a VS200 ASW software (Olympus, Tokyo, Japan) for further analysis.

### Isolation of cells from peripheral blood, bronchoalveolar lavage, and the lung

Cells from peripheral blood, bronchoalveolar lavage (BAL), and lung tissues were isolated as previously described^12^, mice were anesthetized with isoflurane and peripheral blood was exhaustively collected from the abdominal vein. Following exhaustive bronchoalveolar lavage with PBS, lung lobes were harvested, cut into small pieces, and digested in collagenase type I (Thermo Fisher, Waltham, MA) for 1 hour at 37 °C in a shaker incubator. Lung single-cell suspension was obtained by crushing the digested lung tissues through a 100 μm basket filter (BD Biosciences, San Jose, CA). Following erythrocyte lysis of peripheral blood and lung cells by incubation with red blood cell lysing buffer (SolarBio, Beijing, China), cells were washed with PBS and resuspended in either PBS for flow cytometry analysis, or in PBS containing 0.5 % FBS and 2 mM EDTA for magnetic cell purification.

### *In vivo* labeling of AMs and monocytes

AMs were labeled with PKH26 as previously described^16^. Briefly, PKH26 cell labeling kit (MCE) was prepared according to manufacturer’s recommendation and a single dose of PKH26 was administered intranasally into the lungs of mice (10 μM, 50 μL per mouse) at 5 days before IAV infection.

To label peripheral blood monocytes in Cx3cr1-YFP^CreER^R26^tdTomato^ mice, tamoxifen (MCE) dissolved in corn oil (at a final concentration of 15 mg/mL) were injected i.p. in mice with 75 μg tamoxifen per gram body weight per day for 5 consecutive days. Fluorescence intensity of YFP and tdTomato in peripheral blood monocytes was determined by flow cytometry analysis immediately after the final dose of tamoxifen.

### Blockade of IFNAR1 *in vivo*

For *in vivo* blocking of IFNAR1, mice were intranasally administered with a single dose of anti-IFNAR1 monoclonal antibody (BioXcell, clone MAR1-5A3) at 40 μg per mouse diluted in 50 μL of PBS at one day before IAV infection.

### Flow cytometry

Single cell suspension from BAL, the lung, and peripheral blood was plated in U-bottom 96-well plates at ≤ 2 × 10^6^ cells per well in PBS. Cells were stained with Zombie Aqua Fixable Viability Kit (Biolegend, San Diego, CA) according to manufacturer’s instructions. Cells were then washed and incubated with anti-CD16/CD32 (clone 2.4G2, BD Biosciences) in PBS containing 0.5% BSA for 15 min on ice and then stained with fluorochrome-labeled antibodies, including anti-CD45 APC-Cy7 (clone 30-F11, BD Biosciences), anti-CD45.1 FITC(clone A20, Biolegend), anti-CD45.2 PerCP-Cy5.5 (clone 104, Biolegend), anti-CD11b PE-Cy7 (clone M1/70, BD Biosciences), anti-CD11b BV605(clone M1/70, Biolegend), anti-CD11b BUV737 (clone M1/70, eBioscience), anti-CD11c APC (clone N418, Biolegend), anti-CD11c PE-Cy7(clone N418, Biolegend), anti-I-A/I-E Alexa Fluor 700 (clone M5/114.15.2, Biolegend), anti-CD3 V450 (clone 17A2, BD Biosciences), anti-CD3 PE-CF594(clone 17A2, Biolegend), anti-CD45R V450(clone RA3-6B2, BD Bioscience), anti-Ly-6C BV711(clone HK1.4, Biolegend), anti-Ly6G-BV650 (clone 1A8, Biolegend), anti-CD24-BV605 (clone M1/69, Biolegend), anti-CD64-PE (clone X54-5/7.1, BD Biosciences), anti-CD64-PE-Cy7 (clone X54-5/7.1, Biolegend), anti-Siglec-F PE-CF594 (clone E50-2440, BD Biosciences), anti-Siglec-F BV421 (clone S17007L, Biolegend), anti-CD4 PE-Cy7 (clone RM4-5, BD Biosciences), anti- CD8-APC-Cy7 (clone 53-6.7, BD Bioscience), anti-CD119-biotin (clone 2E2, Biolegend) and Streptavidin Percp-Cy5.5(eBioscience).

For intracellular staining of p-STAT1, cells stained with surface markers were fixed and permeabilized with fixation/permeabilization buffer (Thermo Fisher) and washed twice with 1× permeabilization buffer (Thermo Fisher), followed by intracellular staining with anti-p-STAT1 (clone 58D6, CST, 1:200) diluted in permeabilization buffer for 0.5 hour at room temperature. After staining, the cells were washed again and labeled with donkey anti-rabbit IgG AF647 (Invitrogen).

Stained cells were analyzed on a BD LSRFortessa flow cytometer. Data were analyzed and illustrated by using FlowJo software (version 10.7.1; BD Biosciences).

For fluorescence-activated cell sorting of AMs, cells stained with fluorochrome-conjugated antibodies and PKH26 were sorted on a BD FACSAria Fusion cell sorter.

### Parabiotic mouse model

To generate parabiotic mice, the mirroring lateral sides of two age- and weight-matched anesthetized mice (CD45.1^+^ and CD45.2^+^, respectively) were shaved, and skin incisions from elbow to knee joints on the respective mirroring sides were made. The adjacent elbow and knee joints, and the dorsal and ventral skin were respectively approximated by sutures. Carprofen and buprenorphine were administered intraperitoneally on days 0 to 3 of surgery. Three weeks after surgery, both mice were intranasally administered with PBS or IAV as described above. On days 8 and 30 post viral infection, chimerisms in peripheral blood monocytes and alveolar macrophages in BAL and the lung were determined by flow cytometry analysis. Chimerism was respectively calculated as % CD45.1^+^ / (% CD45.1^+^ + % CD45.2^+^) in CD45.2^+^ mice, and as % CD45.2^+^ / (% CD45.2^+^ + % CD45.1^+^) in CD45.1^+^ mice.

### Purification, adoptive transfer, and *ex vivo* stimulation of AMs

Purification, adoptive transfer and *ex vivo* stimulation of AMs were performed as previously described. To obtain purified AMs, BAL cells from naïve or IAV-infected mice were labeled with CD11c microbeads (Miltenyi, Auburn, CA) followed by positive magnetic selection on a MS column magnetic purification system (Miltenyi, Auburn, CA) according to the manufacturer’s instructions. Purified AMs were used for RNA sequencing, ATAC sequencing, adoptive transfer, or *ex vivo* cell culture experiments. For adoptive transfer experiments, pooled AMs from multiple mice from the same treatment group were resuspended at 1×10^6^ AMs per recipient in 50 μL PBS and was administered intratracheally into isoflurane anesthetized mice. For *ex vivo* stimulation, isolated AMs were resuspended in complete RPMI 1640 medium without penicillin/streptomycin (P/S-free media) and plated at 5×10^4^ cells/well in 96-well tissue culture plates. Cells were incubated at 37 °C in a 5 % CO2 incubator for 1 hour prior to stimulation. Live *Streptococcus pneumoniae* were grown to mid-log phase as mentioned above and washed twice in PBS and resuspended in P/S-free media. Bacteria were supplemented to AM culture wells at a MOI = 10 and incubated for 1 hour at 37 °C. After incubation, culture wells were washed twice with PBS followed by supplementing with complete RPMI 1640 containing 1% penicillin/streptomycin and incubation for additional 30 minutes to kill extracellular bacteria. AMs were washed twice with PBS, supplemented with complete RPMI 1640, and cultured for additional 12 hours before the culture supernatants were collected for ELISA assay. Alternatively, AM culture wells were supplemented with LPS at 10 ng/ml and incubated for 12 hours before supernatants were harvested.

### *Ex vivo* bacteria phagocytosis and killing assay

AMs isolated from BAL of naïve or IAV-infected mice were resuspended in P/S-free complete RPMI 1640 medium and were plated at 1×10^5^ cells per well in 96-well tissue culture plates. Cells were incubated at 37 in a 5% CO_2_ incubator for 1 hour prior to stimulation. CFSE labeled live *Streptococcus pneumoniae* were supplemented to AM culture wells at a multiplicity of infection (MOI) = 100 and incubated for 1 hour at 37 . After incubation (denoted as 0h), cells were washed twice with PBS followed by supplementing with complete RPMI 1640 containing 1% P/S and incubation for additional 1.5 hours (denoted as 1.5h). At 0h or 1.5h, AMs were harvested and stained with anti-CD11c APC and anti-SiglecF BV421 antibodies, followed by flow cytometry analysis to determine the CFSE fluorescence in AMs representing bacterial phagocytosis (0h). Bacteria killing was calculated as: % [(MFI of CFSE in AMs at 0h) – (MFI of CFSE in AMs at 1.5h)]/ (MFI of CFSE in AMs at 0h).

### Induction of TI in AMs *ex vivo*

AMs obtained from BAL of naïve wild type C57BL/6 mice were resuspended in complete RPMI 1640 containing 1% penicillin/streptomycin, plated (50,000 cells per well in 96-well cell culture plates) and rested for 2 h at 37 °C in a 5% CO_2_ incubator. For induction of TI *ex vivo*, recombinant IFN-γ (50 ng/ml, Peprotech) and/or IFN-β (500 pg/ml, Peprotech) was supplemented to selected culture wells. After cytokine stimulation (training) for 24 hours, AMs were washed twice with culture medium (without recombinant IFNs) and rested for 6 days. After resting, AMs were re-stimulated with LPS or live S.P. for 12 hours and supernatants were collected for ELISA analysis^12^. Alternatively, *ex vivo* trained AMs were cocultured with B16-luc melanoma cells for 72h to determine tumor cell survival and the concentrations of cytokines in supernatants.

### Coculture of AMs and tumor cells for antitumor activity assay

AMs were isolated as described above and were seeded into a 96-well tissue culture plate at 0.5 × 10^5^ to 2 × 10^5^ cells per well. Cells were incubated in a 37 5% CO_2_ cell-culture incubator for 2 hours. B16-luc or B16-GFP tumor cells were seeded into AM culture wells at 1×10^4^ cells per well. At 24, 48 or 72 hours of coculture, cells in each well was lysed and luciferase activity per well was determined by using Luciferase assay system (Yeason Biotechnology, Shanghai, China) according to manufacturer’s instructions. Luminescence measurements were performed on a SynergyMx M5 plate reader (Molecular Devices, San Jose, CA). Tumor cell survival was calculated and presented as percentage of normalized luminescence against tumor-only culture wells.

### ELISA assay

ELISA assay was performed by using murine TNF, IFN-γ, IL-6, IL-12, and MIP-2 ABTS ELISA development kits (Peprotech, Rocky Hill, NJ) according to manufacturers’ instructions. ELISA plates were read on a SynergyMx M5 plate reader (Molecular Devices, San Jose, CA). Concentrations of cyto-/chemokines were calculated based on serial dilutions of standards by using GraphPad Prism software (Version 9, GraphPad Software, La Jolla, CA).

### Metabolic assay of AMs

Real-time cell metabolism of AMs was determined by using the Seahorse XF Cell Mito Stress Test Kit or Seahorse XF Glycolytic Stress Test Kit (Agilent Technologies, Santa Clara, CA) according to the manufacturer’s instructions. Purified AMs were seeded into XF96-well plates (Agilent Technologies, Santa Clara, CA) at 50,000 cells per well and cultured overnight in complete RPMI 1640 media. On the next day, AMs were washed and incubated for 1 hour in Seahorse assay media (Agilent Technologies, Santa Clara, CA) supplemented with either 2 mM Glutamine (glycolytic stress test) or an additional 10 mM glucose and 1 mM pyruvate (mito stress test). The kit compounds were resuspended in assay media and the final concentrations of the compounds in culture wells are 1.5 μM oligomycin, 4 μM FCCP and 0.5 μM rotenone/antimycin A in mito stress test, or 10 mM glucose, 1 μM oligomycin and 50 mM 2-DG in glycolytic stress test. Oxygen consumption rate (OCR) and extracellular acidification rate (ECAR) were measured on a Seahorse XFe96 Analyzer (Agilent Technologies, Santa Clara, CA) and data were analyzed using Wave Desktop v.2.6 software.

### RNA sequencing and data analysis

A total of 400,000 purified AMs per replicate were pelleted and lysed in TRIzol reagent (Thermo Fisher, Waltham, MA) and cryopreserved at -80 °C, followed by RNA extraction. RNA quantification and qualification were performed using a N50 NanoPhotometer microvolume spectrophotometer (IMPLEN, Westlake Village, CA) and RNA Nano 6000 Assay Kit of the Bioanalyzer 2100 system (Agilent Technologies, Santa Clara, CA). Sequencing libraries were prepared using NEBNext UltraTM RNA Library Prep Kit (New England Biolabs, Ipswich, MA) following manufacturer’s recommendations and index codes were added to attribute sequences to each sample. The clustering of the index-coded samples was performed on a cBot Cluster Generation System using TruSeq PE Cluster Kit v3-cBot-HS (Illumina, San Diego, CA) according to the manufacturer’s instructions. After cluster generation, the library preparations were sequenced on an Illumina Novaseq platform (Illumina, San Diego, CA).

Raw RNA sequencing fastq format data were processed through in-house perl scripts and clean data were obtained by removing reads containing adapter or poly-N and low-quality reads from raw data. Reads were then mapped to mm10 RefSeq genes downloaded from the UCSC Table Browser using Hisat2(v2.0.5). FeatureCounts(v1.5.0-p3) was used to count the reads numbers mapped to each gene. And FPKM (expected number of Fragments Per Kilobase of transcript sequence per Millions base pairs sequenced) of each gene was calculated based on the length and reads count mapped to the gene. Principle component analysis (PCA) plot and differential gene expression analysis was performed by using the DESeq2 (v.1.44.0) package in R program (v.4.4.1). The resulting P-values were adjusted using the Benjamini and Hochberg’s approach for controlling the false discover. An adjusted P value < 0.05 found by DESeq2 was used as the significance threshold for identification of differentially expressed genes. Volcano and heatmap plots were created using the ggplot2 and pheatmap packages separately in R software. Gene ontology (GO) enrichment analyses of differentially expressed genes were performed using the clusterProfiler R package. Gene ontology terms with corrected P value < 0.05 were considered significantly enriched. And clusterProfiler R package was used to test the statistical enrichment of differential expression genes in KEGG pathways. The local version of the GSEA analysis tool http://www.broadinstitute.org/gsea/index.jsp, GO, KEGG data set were used for GSEA independently.

### ATAC sequencing and data analysis

A total of 50,000 purified AMs per replicate were washed once in cold PBS and resuspended in 50 µl of cold lysis buffer (10 mM Tris-HCl, pH 7.4, 10 mM NaCl, 3 mM MgCl2, 0.1% IGEPAL CA-630). Cell lysates were centrifuged at 500g for 10 min at 4 and nuclei were resuspended in 50 µL of transposition reaction mix (25 µL of TD buffer + 2.5 µL of Tn5 Transposase + 22.5 µL of nuclease-free water; Illumina, San Diego, CA) and incubated for 30 min at 37 °C. Transposed DNA fragments were purified by using a Qiagen Reaction MiniElute Kit (Qiagen, Hilden, Germany), barcoded with Nextera single indexes (New England Biolabs, Ipswich, MA), and amplified by PCR for 11 cycles using NEBNext High Fidelity 2× PCR Master Mix (New England Biolabs, Ipswich, MA). PCR products were purified using a PCR Purification Kit (Qiagen, Hilden, Germany) and amplified fragment size was verified on an Agilent 2100 Bioanalyzer system (Agilent, Santa Clara, CA). ATAC-seq libraries were quantified by quantitative PCR. Paired-end sequencing was performed on an Illumina Novaseq6000 system (Novogene, Beijing, China).

Raw ATAC-seq fastq files from paired-end sequencing were trimmed by using fastp (v0.19.11) followed by quality control using fastqc(v0.11.5). Clean data were aligned to the GRCm38/mm10 reference genome using BWA(v0.7.12-r1039). Samtools(v1.11) was used to remove unmapped, unpaired and mitochondrial reads. PCR duplicates were removed by using Picard. Peak calling was performed using MACS2(v2.1.2, -q 0.05 --call-summits --nomodel -- shift -100 --extsize 200 --keep-dup all). For each experiment, peaks of all samples were combined to create a union peak list and overlapping peaks were merged with BedTools merge. The number of reads in each peak was determined using DiffBind(v3.14.0). Differentially accessible peaks were identified following DESeq2 normalization using an FDR cut-off <0.05. Tracks were visualized using Integrative Genomics Viewer (v.2.3.77, Broad Institute). Pearson correlations and PCA between samples were calculated in R(DiffBind v3.14.0) and plotted. Gene ontology term enrichment was performed for each biological condition using clusterProfiler(v4.12.0). Statistical analysis of differential chromatin-accessibility tests was done using DESeq2 (v.1.44.0), and FDR correction was performed using the Benjamini–Hochberg method in R (v.4.4.1). Significance of gene ontology term enrichments was calculated with binomial tests and hypergeometric tests. P values and q values of <0.05 were considered to indicate a significant difference.

### Annalysis of public human scRNA-seq dataset

scRNA-seq data of 10 healthy human adults were downloaded from the Gene Expression Omnibus (GEO) with accession code GSE151928. The Seurat package (version 5.1.0) was used for the data quality control and analysis, as comprehensively outlined by the package developer. Samples were filtered for cell barcodes recording > 500 UMI, with < 20% mitochondrial gene expression. Data were corrected with harmony. Neighbor analysis was performed by invoking FindNeighbors () using the harmony reduction method and the first 30 principle components. Clustering was then performed with FindClusters() at resolution = 0.3 (resulting in 13 clusters). PCA analysis was performed based on the scaled data with the top 2,000 highly variable genes, and the top 30 principals were used for UMAP construction. Differentially expressed genes (DEGs) were identified using the default “FindMarkers” function in Seurat based on the non-parametric Wilcoxon rank–sum test^59^, and analyzed with R program clusterProfiler for GO enrichment analysis. Data from human NSCLC lung tissue scRNA-seq (GSE154826) were analyzed as described in our previous study^12^ and genes of AMs from immune activation high and low groups were used for GSEA enrichment analysis.

### Statistical analysis

Statistical parameters including the exact value of n, the definition of center, dispersion and precision measures, and statistical significance are reported in Figures and Figure Legends. A p value of < 0.05 was considered significant. Two-tailed Student t test was performed for comparisons between two groups. One-way ANOVA followed by a Tukey test was performed to compare more than two groups. All statistical analyses were performed using GraphPad Prism software (Version 9, GraphPad Software, La Jolla, CA).

## References

1. Netea, M. G. et al. Trained immunity: A program of innate immune memory in health and disease. Science 352, aaf1098 (2016).

2. Divangahi, M. et al. Trained immunity, tolerance, priming and differentiation: distinct immunological processes. Nat Immunol 22, 2–6 (2021).

3. Netea, M. G. et al. Defining trained immunity and its role in health and disease. Nat Rev Immunol 20, 375–388 (2020).

4. Cheng, S.-C. et al. mTOR- and HIF-1alpha-mediated aerobic glycolysis as metabolic basis for trained immunity. Science 345, 1250684 (2014).

5. Saeed, S. et al. Epigenetic programming of monocyte-to-macrophage differentiation and trained innate immunity. Science 345, 1251086 (2014).

6. Mulder, W. J. M., Ochando, J., Joosten, L. A. B., Fayad, Z. A. & Netea, M. G. Therapeutic targeting of trained immunity. Nat Rev Drug Discov 18, 553–566 (2019).

7. Wang, T., Wang, Y., Zhang, J. & Yao, Y. Role of trained innate immunity against mucosal cancer. Current Opinion in Virology vol. 64 Preprint at 10.1016/j.coviro.2024.101387 (2024).

8. van Puffelen, J. H. et al. Trained immunity as a molecular mechanism for BCG immunotherapy in bladder cancer. Nat Rev Urol 17, 513–525 (2020).

9. Ding, C. et al. Inducing trained immunity in pro-metastatic macrophages to control tumor metastasis. Nat Immunol 24, 239–254 (2023).

10. Priem, B. et al. Trained Immunity-Promoting Nanobiologic Therapy Suppresses Tumor Growth and Potentiates Checkpoint Inhibition. Cell 183, 786–801.e19 (2020).

11. Yao, Y. et al. Induction of Autonomous Memory Alveolar Macrophages Requires T Cell Help and Is Critical to Trained Immunity. Cell 175, 1634–1650.e17 (2018).

12. Wang, T. et al. Influenza-trained mucosal-resident alveolar macrophages confer long-term antitumor immunity in the lungs. Nat Immunol (2023) doi:10.1038/s41590-023-01428-x.

13. Arts, R. J. W. et al. BCG Vaccination Protects against Experimental Viral Infection in Humans through the Induction of Cytokines Associated with Trained Immunity. Cell Host Microbe 23, 89–100.e5 (2018).

14. Giamarellos-Bourboulis, E. J. et al. Activate: Randomized Clinical Trial of BCG Vaccination against Infection in the Elderly. Cell 183, 315–323.e9 (2020).

15. Kaufmann, E. et al. BCG Educates Hematopoietic Stem Cells to Generate Protective Innate Immunity against Tuberculosis. Cell 172, 176–190.e19 (2018).

16. Jeyanathan, M. et al. Parenteral BCG vaccine induces lung-resident memory macrophages and trained immunity via the gut–lung axis. Nat Immunol 23, 1687–1702 (2022).

17. Mata-Martínez, P., Bergón-Gutiérrez, M. & del Fresno, C. Dectin-1 Signaling Update: New Perspectives for Trained Immunity. Frontiers in Immunology vol. 13 Preprint at 10.3389/fimmu.2022.812148 (2022).

18. Khan, N. et al. M. tuberculosis Reprograms Hematopoietic Stem Cells to Limit Myelopoiesis and Impair Trained Immunity. Cell 183, 752–770.e22 (2020).

19. Murphy, J., Summer, R., Wilson, A. A., Kotton, D. N. & Fine, A. The prolonged life-span of alveolar macrophages. Am J Respir Cell Mol Biol 38, 380–385 (2008).

20. Aegerter, H., Lambrecht, B. N. & Jakubzick, C. V. Biology of lung macrophages in health and disease. Immunity vol. 55 1564–1580 Preprint at 10.1016/j.immuni.2022.08.010 (2022).

21. Kopf, M., Schneider, C. & Nobs, S. P. The development and function of lung-resident macrophages and dendritic cells. Nat Immunol 16, 36–44 (2015).

22. Eguiluz-Gracia, I. et al. Long-term persistence of human donor alveolar macrophages in lung transplant recipients. Thorax 71, 1006–1011 (2016).

23. Zahalka, S. et al. Trained immunity of alveolar macrophages requires metabolic rewiring and type 1 interferon signaling. Mucosal Immunol 15, 896–907 (2022).

24. Leopold Wager, C. M., et al. IFN-gamma immune priming of macrophages in vivo induces prolonged STAT1 binding and protection against Cryptococcus neoformans. PLoS Pathog 14, e1007358 (2018).

25. Lazear, H. M., Schoggins, J. W. & Diamond, M. S. Shared and Distinct Functions of Type I and Type III Interferons. Immunity vol. 50 907–923 Preprint at 10.1016/j.immuni.2019.03.025 (2019).

26. Connolly, E. & Hussell, T. The Impact of Type 1 Interferons on Alveolar Macrophage Tolerance and Implications for Host Susceptibility to Secondary Bacterial Pneumonia. Frontiers in Immunology vol. 11 Preprint at 10.3389/fimmu.2020.00495 (2020).

27. Kamada, R. et al. Interferon stimulation creates chromatin marks and establishes transcriptional memory. Proc Natl Acad Sci U S A 115, E9162–E9171 (2018).

28. Aegerter, H. et al. Influenza-induced monocyte-derived alveolar macrophages confer prolonged antibacterial protection. Nat Immunol 21, 145–157 (2020).

29. Li, F., et al. Monocyte-Derived Alveolar Macrophages Autonomously Determine Severe Outcome of Respiratory Viral Infection. Sci. Immunol vol. 7 https://www.science.org (2022).

30. Lavin, Y. et al. Tissue-resident macrophage enhancer landscapes are shaped by the local microenvironment. Cell 159, 1312–1326 (2014).

31. Gibbings, S. L. et al. Transcriptome analysis highlights the conserved difference between embryonic and postnatal-derived alveolar macrophages. Blood 126, 1357–1366 (2015).

32. Lercher, A. et al. Antiviral innate immune memory in alveolar macrophages after SARS-CoV-2 infection ameliorates secondary disease caused by influenza A virus. Immunity (2024) doi:10.1016/j.immuni.2024.08.018.

33. Yona, S. et al. Fate Mapping Reveals Origins and Dynamics of Monocytes and Tissue Macrophages under Homeostasis. Immunity 38, 79–91 (2013).

34. Zhang, J. et al. Bacterial pneumonia induces senescence in resident alveolar macrophages that are outcompeted by monocytes. Cell Rep 44, (2025).

35. Mizutani, T. et al. Conditional IFNAR1 ablation reveals distinct requirements of Type I IFN signaling for NK cell maturation and tumor surveillance. Oncoimmunology 1, 1027– 1037 (2012).

36. Ivashkiv, L. B. IFNγ: signalling, epigenetics and roles in immunity, metabolism, disease and cancer immunotherapy. Nature Reviews Immunology vol. 18 545–558 Preprint at 10.1038/s41577-018-0029-z (2018).

37. Novakovic, B. et al. β-Glucan Reverses the Epigenetic State of LPS-Induced Immunological Tolerance. Cell 167, 1354–1368.e14 (2016).

38. Mould, K. J. et al. Airspace macrophages and monocytes exist in transcriptionally distinct subsets in healthy adults. Am J Respir Crit Care Med 203, 946–956 (2021).

39. Leader, A. M. et al. Single-cell analysis of human non-small cell lung cancer lesions refines tumor classification and patient stratification. Cancer Cell 39, 1594–1609.e12 (2021).

40. Singh, A. K. et al. Re-engineered BCG overexpressing cyclic di-AMP augments trained immunity and exhibits improved efficacy against bladder cancer. Nat Commun 13, (2022).

41. Netea, M. G. et al. Trained Immunity: a Tool for Reducing Susceptibility to and the Severity of SARS-CoV-2 Infection. Cell 181, 969–977 (2020).

42. Joseph, J. Trained Immunity as a Prospective Tool against Emerging Respiratory Pathogens. Vaccines (Basel) 10, 1932 (2022).

43. Netea, M. G., Joosten, L. A. B. & van der Meer, J. W. M. Hypothesis: stimulation of trained immunity as adjunctive immunotherapy in cancer. J Leukoc Biol 102, 1323–1332 (2017).

44. Lérias, J. R. et al. Trained Immunity for Personalized Cancer Immunotherapy: Current Knowledge and Future Opportunities. Front Microbiol 10, 2924 (2019).

45. Geller, A. E. et al. The induction of peripheral trained immunity in the pancreas incites anti-tumor activity to control pancreatic cancer progression. Nat Commun 13, (2022).

46. Van Puffelen, J. H. et al. Intravesical BCG in patients with non-muscle invasive bladder cancer induces trained immunity and decreases respiratory infections. J Immunother Cancer 11, (2023).

47. Kalafati, L. et al. Innate Immune Training of Granulopoiesis Promotes Anti-tumor Activity. Cell 183, 771–785.e12 (2020).

48. Zitvogel, L., Galluzzi, L., Kepp, O., Smyth, M. J. & Kroemer, G. Type I interferons in anticancer immunity. Nature Reviews Immunology vol. 15 405–414 Preprint at 10.1038/nri3845 (2015).

49. Holicek, P., et al. Type I interferon and cancer. Immunological Reviews vol. 321 115–127 Preprint at 10.1111/imr.13272 (2024).

50. Swiecki, M. & Colonna, M. Type I interferons: diversity of sources, production pathways and effects on immune responses. Curr Opin Virol 1, 463–475 (2011).

51. Chin, E. N., Sulpizio, A. & Lairson, L. L. Targeting STING to promote antitumor immunity. Trends in Cell Biology vol. 33 189–203 Preprint at 10.1016/j.tcb.2022.06.010 (2023).

52. Schliehe, C. et al. The methyltransferase Setdb2 mediates virus-induced susceptibility to bacterial superinfection. Nat Immunol 16, 67–74 (2015).

53. Sun, L. et al. Activating a collaborative innate-adaptive immune response to control metastasis. Cancer Cell 39, 1361–1374.e9 (2021).

54. Careau, E. & Bissonnette, E. Y. Adoptive transfer of alveolar macrophages abrogates bronchial hyperresponsiveness. Am J Respir Cell Mol Biol 31, 22–27 (2004).

55. Suzuki, T. et al. Pulmonary macrophage transplantation therapy. Nature 514, 450–454 (2014).

56. Casanova-Acebes, M. et al. Tissue-resident macrophages provide a pro-tumorigenic niche to early NSCLC cells. Nature 595, 578–584 (2021).

57. Sharma, S. K. et al. Pulmonary alveolar macrophages contribute to the premetastatic niche by suppressing antitumor T cell responses in the lungs. J Immunol 194, 5529–5538 (2015).

58. Subramanian, S. et al. Long-term culture-expanded alveolar macrophages restore their full epigenetic identity after transfer in vivo. Nat Immunol 23, 458–468 (2022).

59. Hao, Y. et al. Dictionary learning for integrative, multimodal and scalable single-cell analysis. Nat Biotechnol 42, 293–304 (2024).

