## Supplementary figures and legends for "Type-I interferon priming signal balances anti-bacterial and anti-tumor trained immunity in alveolar macrophages"

**Supplementary figures and figure legends**

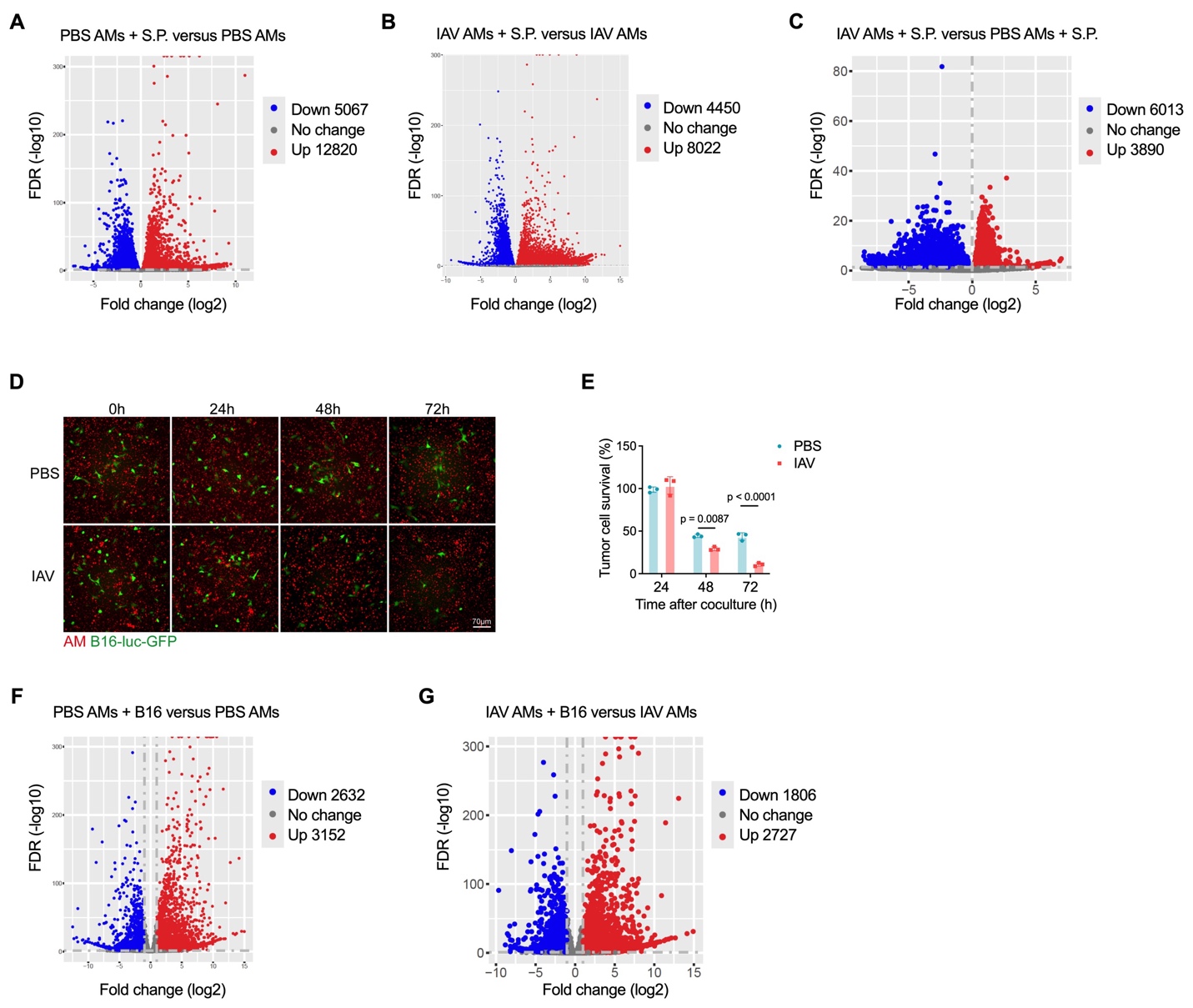

**Figure S1. IAV-trained AMs confer enhanced response to both bacteria and tumor cells (Related to Figure 1)**

(A) RNA-seq volcano plot of differentially expressed genes in AMs isolated from uninfected mice (PBS AMs) with or without ex vivo stimulation with live *Streptococcal pneumoniae* (S.P.).

(B) RNA-seq volcano plot of differentially expressed genes in S.P.-stimulated versus unstimulated AMs isolated from day 30 IAV-infected mice (IAV AMs).

(C) RNA-seq volcano plot of differentially expressed genes in S.P.-stimulated PBS or IAV AMs.

(D) Representative microscopic images on GFP-expressing B16 melanoma cells cocultured with AMs isolated from PBS or day 30 IAV-infected mice and labeled with fluorescence dye. Cells were cocultured for 24, 48 or 72 hours at an AM:B16 (E:T) ratio of 10:1.

(E) Survival of B16 melanoma cells at 24, 48 and 72 hours after coculture with PBS or IAV AMs at an E:T ratio of 10:1.

(F) RNA-seq volcano plot of differentially expressed genes in B16 tumor cell-stimulated versus unstimulated AMs isolated from uninfected mice (PBS AMs).

(G) RNA-seq volcano plot of differentially expressed genes in B16 tumor cell-stimulated versus unstimulated AMs isolated from day 30 IAV-infected mice (IAV AMs).

Bar graphs are presented as mean ± SD. Data in **A**, **B**, **C**, **F**, and **G** are from one experiment with n = 3 mice per group. Data in **D** and **E** are representatives of three independent experiments with n = 3 duplicated culture wells per group. Two-tailed Student t test was performed for comparisons between two groups.

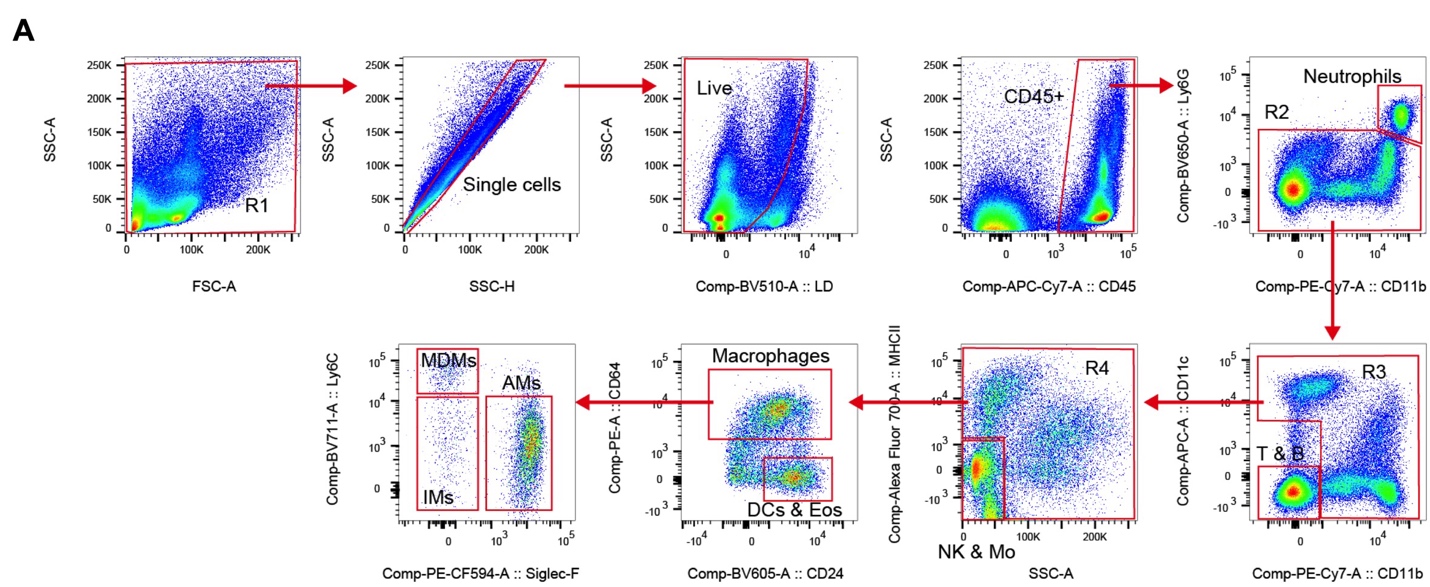

**Figure S2. Flow cytometry gating strategy of immune cell subsets in the lung (Related to Figure 3)**

(A) Representative flow cytometry dot plots showing gating strategy of leukocyte subsets including neutrophils, T and B lymphocytes, NK cells, monocytes (Mo), dendritic cells (DCs), eosinophils (Eos), alveolar macrophages (AMs), interstitial macrophages (IMs), and monocyte-derived macrophages (MDMs), in lung tissues of IAV-infected mice.

Data are representatives of three independent experiments with n = 3 mice per group.

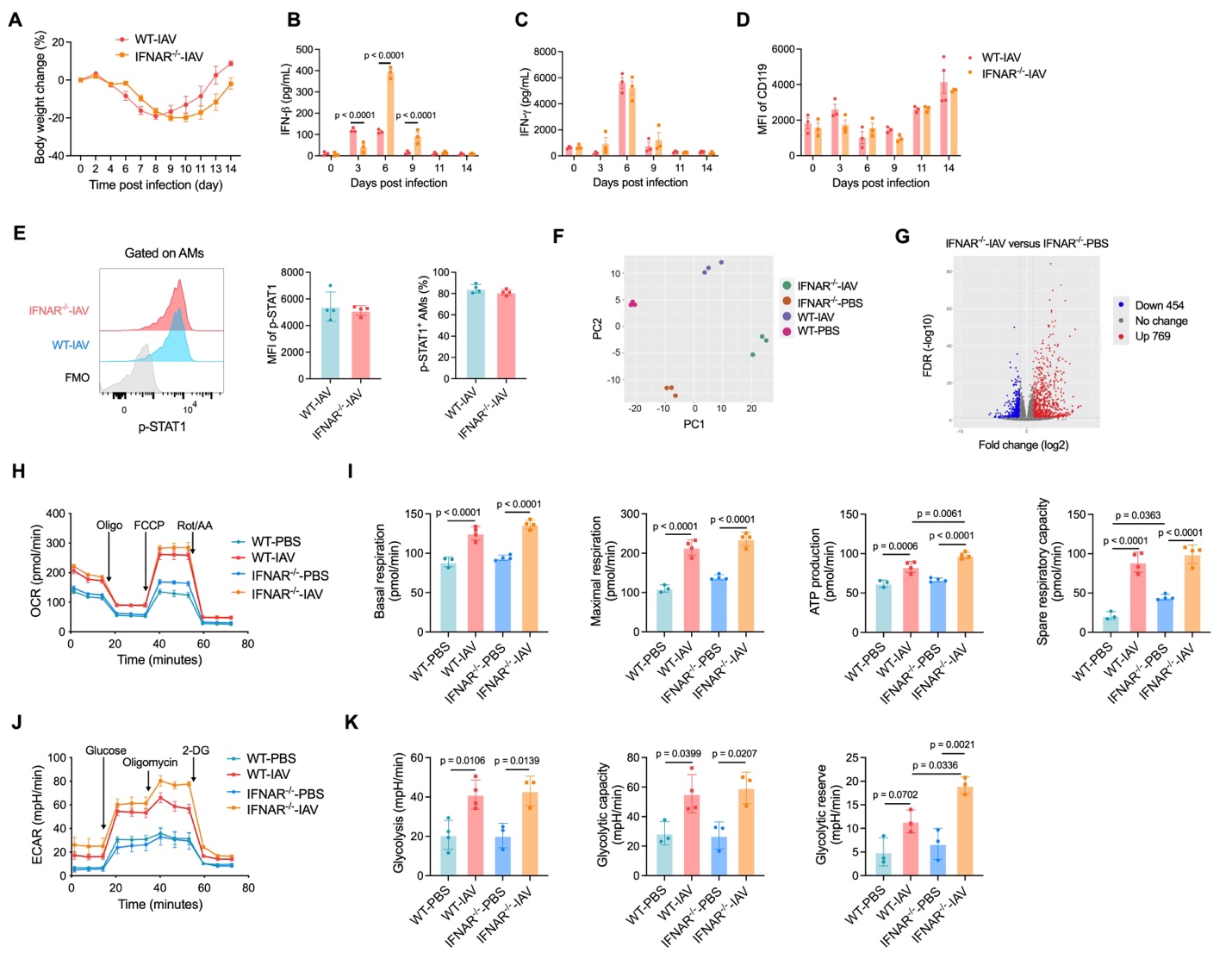

**Figure S3. IAV-infection in IFNAR^-/-^ mice induces TI in AMs (Related to Figure 3)**

(A) Kinetic body weight change in IAV-infected WT or IFNAR^-/-^ mice.

(B,C) Kinetic changes in IFN-β (B) and IFN-γ (C) contents in BAL supernatants post IAV infection.

(D) Median fluorescence intensity (MFI) of CD119 in AMs post IAV infection.

(E) Representative flow cytometry histograms and MFI of intracellular phosphorylated STAT1 (p-STAT1) in AMs, and percentage of p-STAT1 positive AMs, in IAV-infected WT or IFNAR^-/-^ mice at 6 days post infection. FMO, fluorescence minus one.

(F) PCA plot of transcriptional changes of RNA-seq data in AMs from IAV-infected or uninfected WT or IFNAR^-/-^ mice.

(G) Volcano plot of differentially expressed genes in AMs from IAV-infected versus uninfected IFNAR^-/-^ mice.

(H) Real-time oxygen consumption rate (OCR) in AMs from IAV-infected or uninfected WT or IFNAR^-/-^ mice.

(I) Basal respiration, maximal respiration, ATP production and spare respiratory capacity in AMs shown in **H**.

(J) Real-time extracellular acidification rate (ECAR) in AMs.

(K) Glycolysis, glycolytic capacity and glycolytic reserve in AMs shown in **J**.

Graphs with error bars are presented as mean ± SD. Data in **A**-**E** and **H**-**K** are representatives of three independent experiments with n = 3 or 4 duplicated culture wells per group as indicated. Data in **F** and **G** are from one experiment with n = 3 biological replicates per group. Two-tailed Student t test was performed for comparisons between two groups. One-way ANOVA followed by a Tukey test was performed to compare more than two groups.

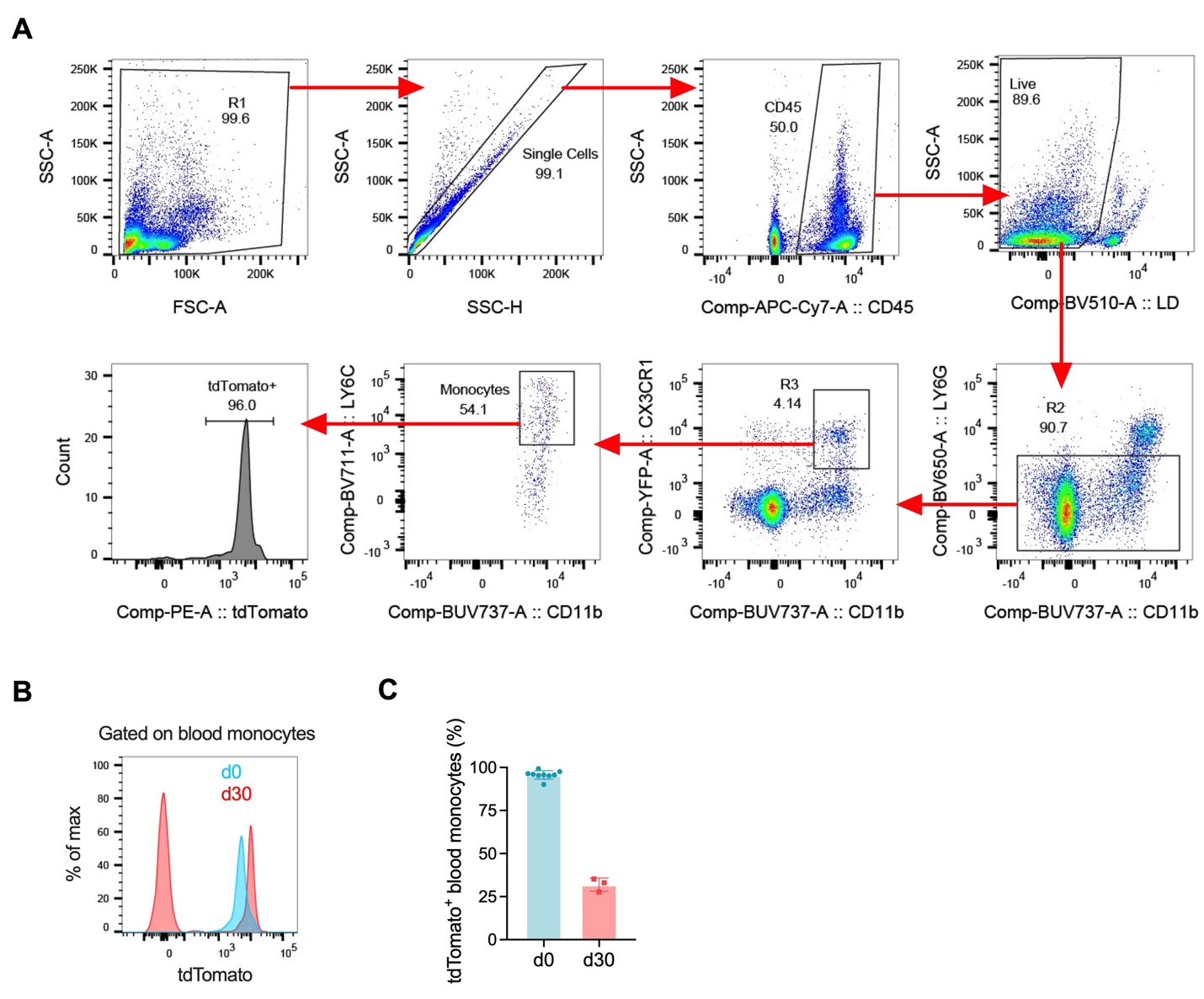

**Figure S4. Tamoxifen induces tdTomato fluorescence in circulating monocytes in Cx3cr1-YFP^CreER^R26^tdTomato^ mice (related to Figure 4)**

(A) Representative flow cytometry dot plots showing gating strategy of circulating monocytes in Cx3cr1-YFP^CreER^R26^tdTomato^ mice after repeated doses of tamoxifen.

(B) Representative flow cytometry histograms of tdTomato fluorescence in peripheral blood monocytes in mice before (d0) and at 30 days (d30) after IAV infection.

(C) Percentages of tdTomato^+^ peripheral blood monocytes before (d0) and at 30 days (d30) after IAV infection.

Data are representatives of two independent experiments with n = 3 mice in IAV-infected group and n = 9 mice in uninfected group.

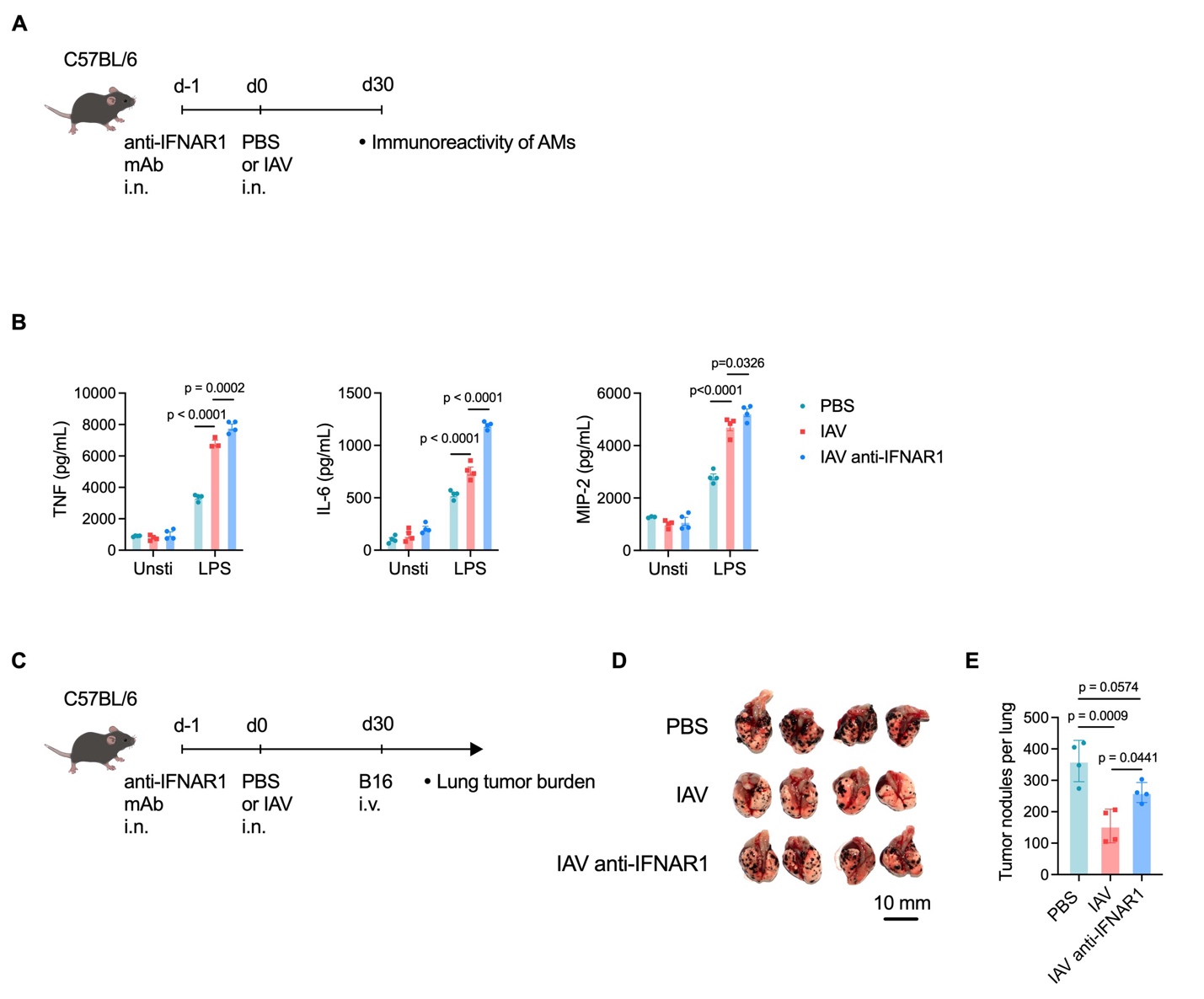

**Figure S5. IFN-I priming signal is required to developing anti-tumor TI functions in AMs (Related to Figure 5)**

(A) Schema of i.n. administration of anti-IFNAR1 monoclonal antibody (mAb) one day before IAV infection in WT mice followed IAV infection, followed by phenotypic and functional analysis of AMs at 30 days post infection.

(B) Concentrations of representative proinflammatory cyto-/chemokines, including TNF, IL-6 and MIP-2, in supernatants of cultured AMs isolated from mice shown in **A**. AMs were either unstimulated (Unsti) or stimulated *ex vivo* with LPS.

(C) Schema of i.n. administration of anti-IFNAR1 mAb one day before IAV infection in WT mice followed by i.v. inoculation of B16 melanoma cells at 30 days post infection. Lung tumor burdens were determined at experimental endpoint.

(D, E) Macroscopic evaluation of the lungs (D) and number of macroscopically visible B16 tumor nodules on the surface of lung lobes in mice showed in **C** (E).

Bar graphs are presented as mean ± SD. Data are representatives of two independent experiments with n = 3 or 4 mice or duplicated culture wells per group as indicated. One-way ANOVA followed by a Tukey test was performed to compare more than two groups.

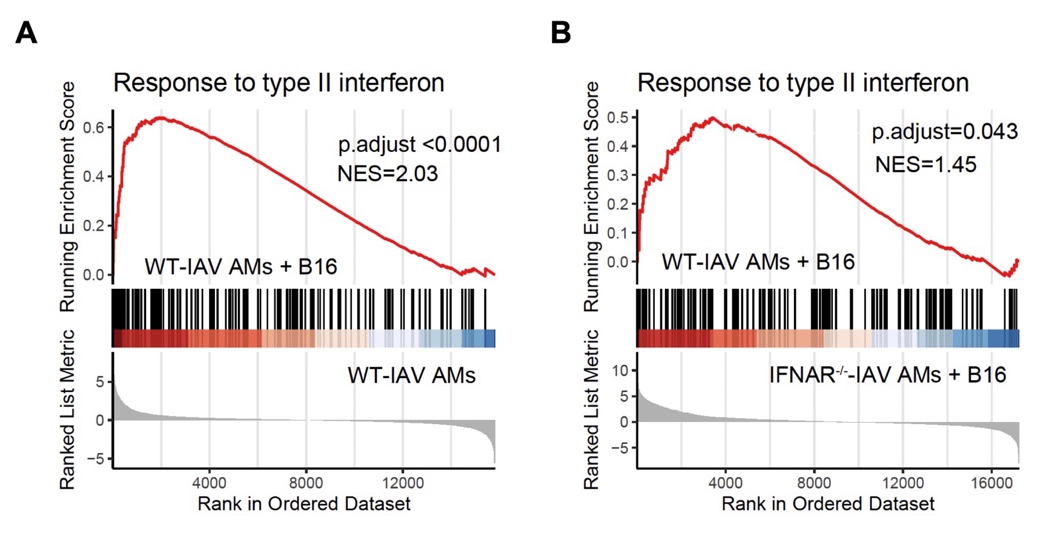

**Figure S6. IFN-I signaling is required to enhanced response to IFN-γ by IAV-trained AMs in response to tumor cells (Related to Figure 6)**

(A) GSEA enrichment of gene transcripts related to response to type II interferon in WT IAV-trained AMs stimulated with B16 tumor cells versus those without tumor cell stimulation.

(B) GSEA enrichment of gene transcripts related to response to type II interferon in B16 melanoma cell-stimulated WT versus IFNAR^-/-^ IAV-trained AMs.

Data are from one experiment with n = 3 biological replicates per group in groups including WT-IAV AMs and WT-IAV AMs + B16, and n = 2 biological replicates per group in IFNAR^-/-^-IAV AMs + B16 group.

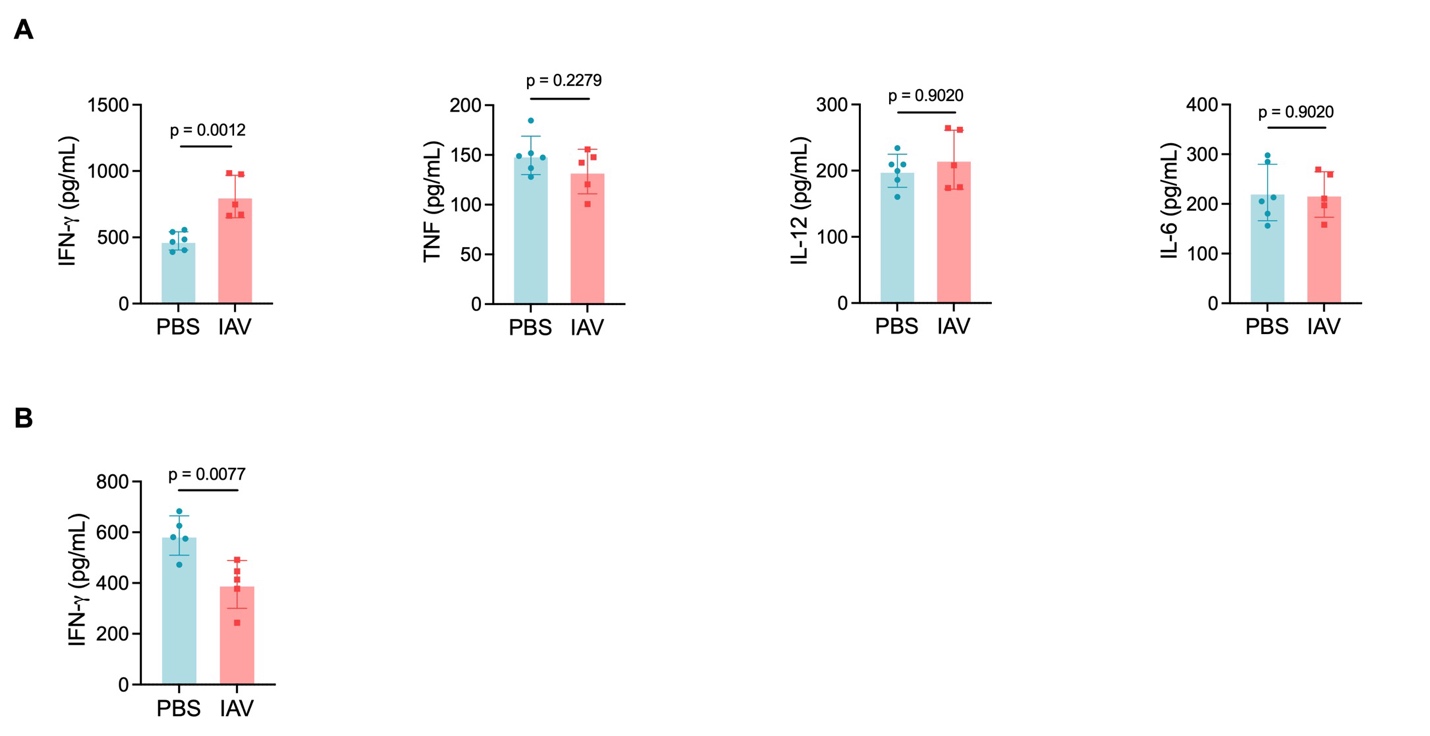

**Figure S7. Prior IAV infection induces increased contents of IFN-γ in lung tissues of tumor-bearing but not S.P.-infected mice (Related to Figure 6)**

(A) Contents of IFN-γ, TNF, IL-12 and IL-6 in the BAL fluid of lung B16 melanoma tumor-bearing

WT mice at 16 days post tumor inoculation. Mice were prior infected with IAV or uninfected (PBS) at 30 days before tumor inoculation.

(B) Contents of IFN-γ in the BAL fluid of mice at 48 hours post respiratory S.P. infection.

Data are representatives of two independent experiments with n = 4, 5, or 6 mice per group as indicated. Two-tailed Student t test was performed for comparisons between two groups.

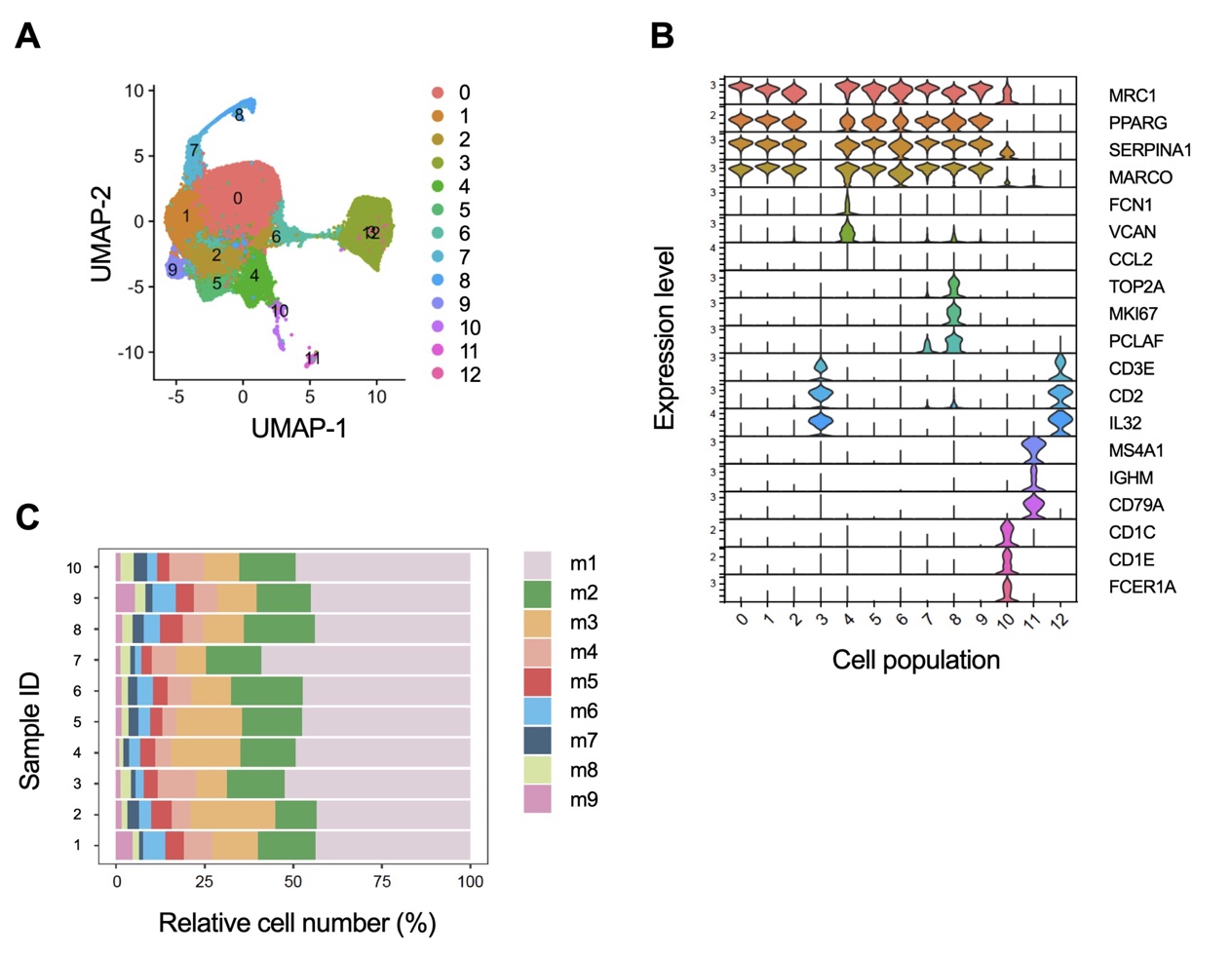

**Figure S8. Subpopulations of human AMs carry transcriptional imprints of IFN-I signaling (Related to Figure 8)**

(A) scRNA-seq UMAP of 13 leukocyte clusters in BAL fluids from healthy human donors.

(B) Volin plot of marker gene transcripts in leukocyte clusters as shown in **A**.

(C) Relative cell number of subpopulations of AMs (m1-m9) in BAL samples (samples ID numbers 1-10).
